# Comparative Analysis of Commercial Single-Cell RNA Sequencing Technologies

**DOI:** 10.1101/2024.06.18.599579

**Authors:** Marco De Simone, Jonathan Hoover, Julia Lau, Hayley Bennet, Bing Wu, Cynthia Chen, Hari Menon, Amelia Au-Yeung, Sean Lear, Samir Vaidya, Minyi Shi, Jessica M. Lund, Ana Xavier-Magalhaes, Yuxin Liang, Ahmet Kurdoglu, William E. O’Gorman, Zora Modrusan, Daniel Le, Spyros Darmanis

## Abstract

This study evaluates ten commercially available single-cell RNA sequencing (scRNA-seq) approaches across four technology groups: Emulsion-based kits from 10x Genomics and Fluent Biosciences; Microwell-based kits from Becton Dickinson, Honeycomb Technologies and Singerlon Technologies; Combinatorial-indexing kits from Parse Biosciences and Scale Biosciences; and a Matrigel-based kit from Scipio Biosciences. Peripheral blood mononuclear cells (PBMCs) from a single donor were used to assess analytical performance. Key features such as sample compatibility, cost, and experimental duration were also compared. Notably, superior analytical performance was demonstrated by the Chromium Fixed RNA Profiling kit from 10x Genomics, which uniquely features probe hybridization for transcript detection. Additionally, the Rhapsody WTA kit from Becton Dickinson provided a cost-effective balance of performance and expense per cell. With a rich dataset of 218,154 cells, this work provides a basis for differentiating commercial scRNA-seq technologies, which is intended to facilitate the effective application and further methodological development of single cell transcriptomics.

## Introduction

Over the past ten years, the emerging landscape of single-cell technologies has ushered in a new era in biomedical research, unveiling previously uncharted terrains of cellular types, states, and lineages ^1^. Pioneering technologies have increased the number of cells retrieved from a single experiment ^2^ while continuously reducing the cost per cell. The initial surge in throughput primarily resulted from the development of droplet-based ^3,4^ and microwell-based methodologies ^5,6^, which have facilitated the parallel processing of thousands of cells. More recently, the advent of combinatorial indexing, in which fixed cells serve as reaction chambers for sequential RNA barcoding, has further increased cell throughput. Combinatorial indexing removes the need to physically isolate each cell, as each cDNA molecule receives a unique combinatorial barcode that distinguishes its cellular origin and enables simultaneous processing of hundreds of thousands of cells ^7,8^.

The commercialization of scRNA-seq technologies, often originating from academic prototypes, has increased access to these techniques. However, given the diverse range of available technologies, a question arises: How do these different strategies compare in terms of performance and effectiveness? Although several evaluations of the available scRNA-seq approaches exist ^9,10,11,12,13^, there currently lacks a systematic evaluation of existing and emerging commercial technologies. This knowledge gap hinders our ability to select an optimal scRNA-seq approach tailored to specific investigative needs. By assessing the performance of each technology at the library, gene, and cell levels, we aim to offer guidance in the application of these tools to biomedical research.

We categorized the methods compared in this study into four general technology groups: emulsion-based methods, microwell-based methods, combinatorial indexing methods and one matrigel-based method **(Fig. 1a)**. To assess performance of the different single-cell RNAseq kits, we used PBMCs from a single donor blood draw. The cell type distribution of this PMBC sample was determined using CyTOF^14^. The PBMCs were split into aliquots and used for each kit assessment. For most kits, two independent technical replicates were performed. However, due to initial cell calling variability, we included a third replicate for Fluent. For Parse and Scale, only a single replicate was processed due to high kit costs.

**Figure 1.**
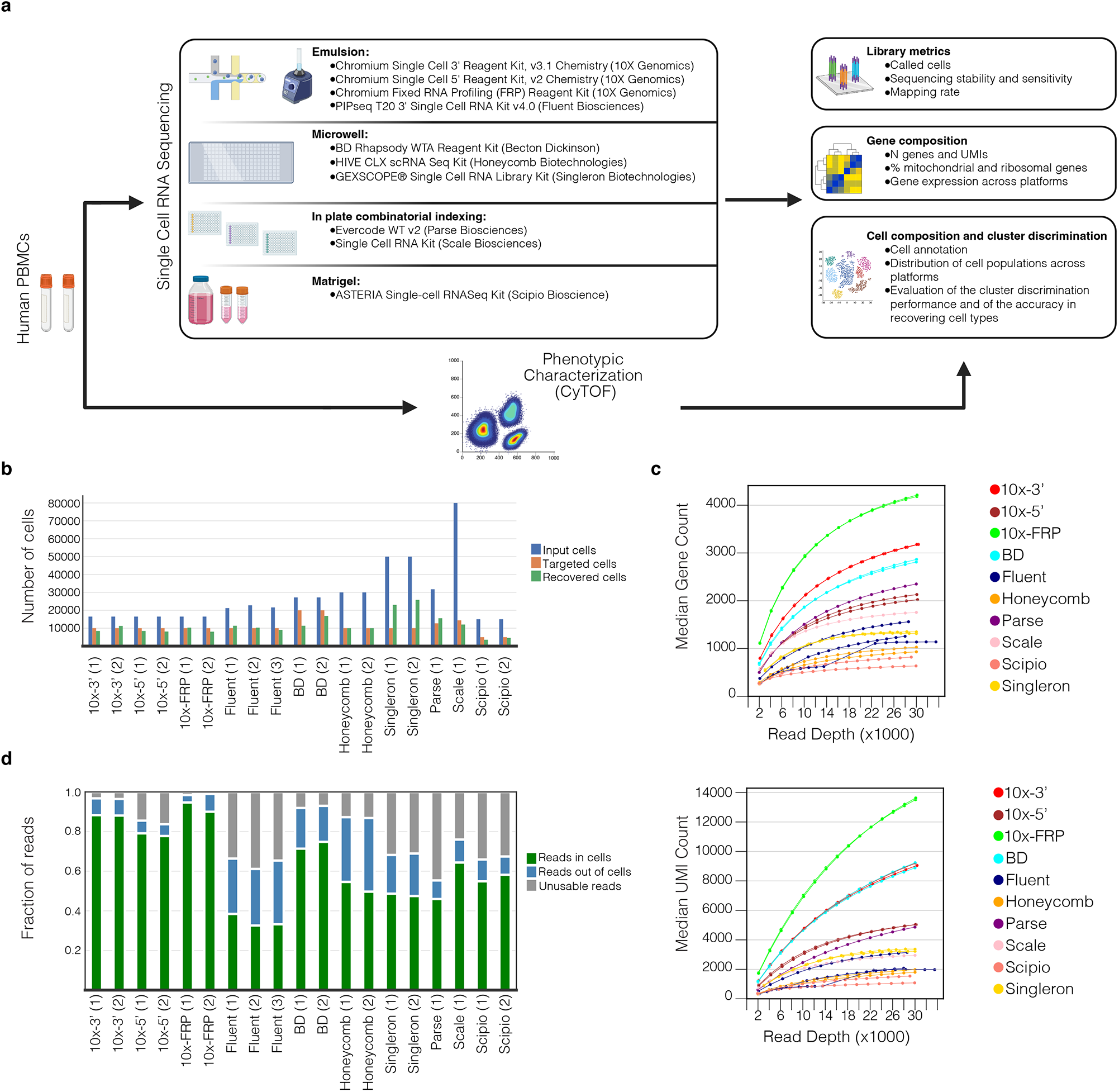
Experimental workflow and library-level metrics. **a)** Experimental layout. Aliquots of the same PBMC sample were processed using ten different commercially available scRNA-seq kits. The same sample was phenotypically characterized using CyTOF. Following library preparation and sequencing, the protocols were assessed for their performance on the library, gene and cell composition level. **b)** Number of input cells (*i.e.* cells loaded on device) used to yield the desired cell target (blue), number of targeted cells (orange), and number of cells recovered by each kit according to the associated cell calling algorithm (green). **c)** Number of median genes (top) and UMIs (bottom) detected at downsampled depths from 2,000 to 30,000 reads per cell in steps of 2,000. Downsampled data were aggregated by kit and fitted with a Michaelis-Menten curve to model saturation. **d)** Read allocation per kit with: (green) the fraction of reads mapped and tagged with a cell barcode, (blue) the fraction of reads mapped and tagged with a non-cell barcode, and (gray) the fraction of unusable reads not mapped (*e.g.* read duplicates) and/or tagged with a cell barcode. Numbers in parentheses denote replicates for each method.

For emulsion-based kits, individual cells are encapsulated in water-in-oil emulsion droplets that are used as reaction chambers. We tested the Chromium Single Cell 3’ reagent kit v3.1 (10x 3’), the Chromium Single Cell 5’ reagent kit v2 (10x 5’), the Chromium Fixed RNA Profiling kit by 10x Genomics (10x FRP), and the PIP Seq T20 3’ Single Cell RNA kit v4 by Fluent Biosciences (Fluent). The 10x 3’, 10x 5’, and Fluent kits utilize Reverse Transcription (RT) to generate barcoded cDNA molecules from mRNA. As the name implies, the 10x 5’ kit targets the 5’-end of cDNA for sequencing. Unique among these kits is the 10x FRP kit, which utilizes the oligonucleotide ligation assay (OLA)^15^ to measure gene expression in formaldehyde-fixed samples using paired probes. Currently, only mouse or human probe sets are available. Cognate probe pairs were designed to hybridize proximally at target loci, enabling probe ligation and capture in droplets (**Extended Data Fig. 1a**).

With microwell-based kits, cells are separated in microwells that serve as individual reaction chambers for barcoding. We tested the BD Rhapsody WTA reagent kit by Becton Dickinson (BD), the HIVE CLX scRNA seq v1 kit by Honeycomb Technologies (Honeycomb), and the Gexscope Single-Cell RNA Library kit by Singleron Technologies (Singleron). Each usable microwell contains a single cell and a magnetic bead conjugated with barcoded oligos. Following cell lysis, mRNA capture and RT yield barcoded cDNA strands that are pooled from all microwells on the device for subsequent pooled second strand synthesis and amplification. BD and Honeycomb use random primers for second strand synthesis while Singleron employs a template switching mechanism (**Extended Data Fig. 1b**).

Combinatorial indexing methods are based on split-pool indexing of cDNA within fixed cells. Through iterative PCR or ligation, a unique set of barcodes is appended to the collection of cDNA molecules from each cell. We tested two combinatorial indexing kits: Evercode WT kit v2 by Parse Biosciences (Parse) and the Single Cell RNA kit by Scale Biosciences (Scale). For both kits, cells are fixed and permeabilized, which facilitates intracellular split-pool indexing without the need to isolate individual cells (**Extended Data Fig. 1c)**. For Parse, barcodes are added through three rounds of ligation and one round of PCR. The Scale kit follows a similar procedure but only requires two rounds of ligation. Additionally, the Parse kit achieves broad transcript coverage using a mix of barcoded random and oligo-dT primers, whereas the Scale kit covers only the 3’-end of transcripts, since it relies solely on oligo-dT priming.

Lastly, we tested the Asteria Single Cell RNA seq kit by Scipio Biosciences (Scipio), which is a method that isolates individual cells paired with barcoded capture beads within a matrigel matrix. After cell lysis, the mRNA content in each cell is hybridized to bead-conjugated oligo-dT sequences containing a unique cell barcode (CB). Then, the matrigel is dissolved to collect bead-bound mRNA that subsequently undergoes RT and cDNA amplification (**Extended Data Fig. 1d**).

## Results

### Library metrics

For each evaluated kit, PBMC aliquots were processed according to the user manual. Generally, each device was loaded such that the expected recovery was 10,000 cells (**Table 1**). Due to the maximum cell yield for the Scipio kit, only 5,000 cells were targeted. For the BD kit, preliminary unpublished results indicated cell recovery around 50% of projected yield; therefore, we targeted 20,000 cells with the expectation of recovering the desired 10,000 cells. The number of recovered cells was determined using companion cell calling algorithms provided with each kit’s software (**Fig. 1b**). Notably, the cell calling algorithms for both Honeycomb and Fluent require user-defined parameter selection that substantially impacts observed recovery. Mean cell recovery based on the number of targeted cells ranged from approximately 70% (BD) to 244% (Singleron) (**Supplementary Table 1-2)**. Consistent with aforementioned preliminary results, BD demonstrated the lowest cell recovery, with one replicate yielding 43% fewer cells than expected according to loading recommendations. To investigate the effect of sequencing depth on cell recovery, we downsampled individual data sets to yield sub-libraries spanning 2,000 to 30,000 reads per cell. Surprisingly, Fluent and Singleron exhibited substantially higher cell recovery variation compared to other kits (**Extended Data Fig. 2a,b**). The results for Singleron were particularly concerning given the linear correlation (mean Pearson r = 0.998) between observed cell count yield and sequencing depth across the downsampling range. This unexpected linear dependence could be solved using an independent cell caller; however, observed cell recovery was still up to 2-fold higher than expected (**Extended Data Fig. 2c)**. These observations call into question the robustness and accuracy of cell calling by Singleron; therefore, Singleron was excluded from subsequent cell-based comparisons.

**Table 1.** Table of kit descriptions and requirements.

| Protocols Tested | Abbreviation | Technology Group | Detection Method | Transcript Coverage | Fixation | Sequencing Format [R1,R2,I1,I2] (bp) | Required Cell Loading |
| --- | --- | --- | --- | --- | --- | --- | --- |
| Chromium Single Cell 3' Reagent Kit, v3.1 Chemistry. (10x Genomics) | 10X 3' | Emulsion | Reverse Transcription | 3-prime end | no | [28,90,10,10] | 16,000 |
| Chromium Single Cell 5' Reagent Kit, v2 Chemistry. (10x Genomics) | 10X 5' | Emulsion | Reverse Transcription | 5-prime end | no | [28,90,10,10] | 16,000 |
| Chromium Fixed RNA Profiling Reagent Kit. (10x Genomics) | 10X FRP | Emulsion | Probe Hybridization | N/A | yes (CHO) | [28,90,10,10] | 16,000 |
| PIPseq T20 3' Single Cell RNA Kit v4.0. (Fluent Biosciences) | Fluent | Emulsion | Reverse Transcription | 3-prime end | no | [54,67,8,8] | 40,000 |
| BD Rhapsody WTA Reagent Kit. (Becton Dickinson) | BD | Microwell | Reverse Transcription | 3-prime end | no | [75,75,8,0] | 30,000 |
| HIVE CLX scRNA Seq Kit. (Honeycomb Biotechnologies) | Honeycomb | Microwell | Reverse Transcription | uniform | no | [25,50,8,8] | 60,000 |
| GEXSCOPE® Single Cell RNA Library Kit. (Singleron Biotechnologies) | Singleron | Microwell | Reverse Transcription | 3-prime end | no | [150,150,8,0] | 50,000 |
| Evercode WT v2. (Parse Biosciences) | Parse | Combinatorial Indexing | Reverse Transcription | uniform | yes (unknown) | [74,86,6,0] | 350,000 |
| Single Cell RNA Kit. (Scale Biosciences) | Scale | Combinatorial Indexing | Reverse Transcription | 3-prime end | yes (MeOH) | [34,76,10,0] | 960,000 |
| ASTERIA Single-cell RNASeq Kit. (Scipio Bioscience) | Scipio | Matrigel | Reverse Transcription | 3-prime end | no | [28,75,6,0] | 15,000 |

Using the downsampled sub-libraries aggregated by kit, we generated total gene and UMI counts at each sequencing depth (**Fig. 1c**). These data were fitted using the Michaelis-Menten saturation model to estimate the maximum number of identifiable genes and UMIs and the number of reads required to reach half-maximal saturation. Hierarchical clustering of the fitted curves indicates three performance tiers based on median detected genes: *Tier 1)* 10x FRP, *Tier 2)* 10x 3’, 10x 5’, BD, Parse, and Scale, and *Tier 3)* Fluent, Singleron, Honeycomb, and Scipio (**Extended Data Fig. 2d,e)**. For median recovered UMIs, we observed two performance tiers with 10x FRP, 10x 3’, and BD performing better than the remaining kits. For both gene and UMI counts, 10x FRP followed by 10x 3’ and BD exhibited the steepest saturation curves and reached the highest number of attainable genes and UMIs (**Supplementary Tables 2-3**).

Using the 30,000 reads per cell sub-library, we analyzed the proportion of reads that were mapped, CB-tagged, and counted toward gene expression (**Fig. 1d)**. The mean proportion of usable reads that were both mapped and associated with a called cell ranged from 34.8% (Fluent) to 92.4% (10x FRP) (**Supplementary Table 4)**. Next, we calculated the average UMI recovery (total UMI counts / total fastq reads) and identified three tiers: *Tier 1)* 10x FRP, *Tier 2)* 10x 3’ and BD, and *Tier 3)* 10x 5’, Parse, Singleron, Scale, Fluent, Honeycomb, and Scipio (**Extended Data Fig. 3a,b,c)**. These tiers differentiate kits into groupings based on the relative utilization of reads, with mean fractions ranging from 0.05 (Scipio) to 0.44 (10x FRP), which ultimately contribute to gene expression counts (**Supplementary Table 5**)

### Gene composition

Next, we examined the number of detected genes and UMIs per cell derived from each kit using count-matched subsamples of cells (n = 7750, excluding Scipio; Scipio-rep-1: n=3457, Scipio-rep2: n=4425) passing quality controls for both metrics (**Fig. 2a**, **Supplementary Table 6**). Hierarchical clustering of median gene and UMI counts per cell revealed two tiers with 10x FRP, 10x 3’ and BD yielding significantly higher gene and UMI counts than the remaining kits (Kruskal-Wallis P value < 0.01, post-hoc Dunn’s test BF-adjusted P value < 0.01) **(Extended Data Fig. 4a, Supplementary Tables 6-8**). Tier 1 kits exhibited median gene counts per cell ranging from 2,793-4,312, while Tier 2 kits displayed a substantially lower range of 627-2,324. Likewise, Tier 1 median UMI counts per cell ranged from 8,779-13,719, which was markedly higher than the 1,063-4,994 range demonstrated by the Tier 2 kits.

**Figure 2.**
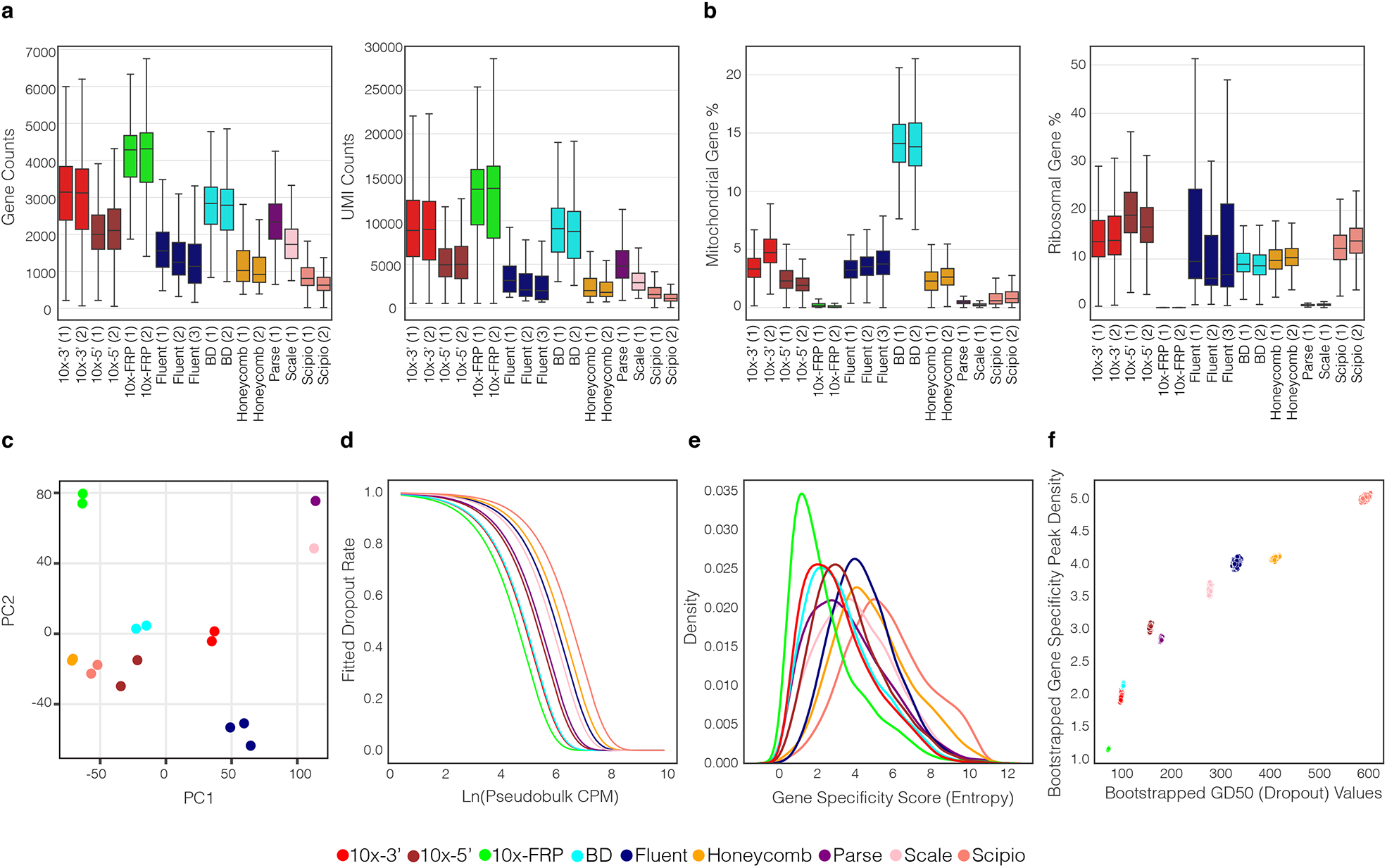
Gene-level metrics. **a)** The distribution of genes (left) and UMIs (right) recovered for each kit. **b)** The distribution of MT gene (left) and RP gene (right) percentages for each kit. **c)** PCA embeddings based on pseudobulk mean gene expression. Numbers in parentheses denote replicates for each method. **d)** Exponential decay fitted values of mean bootstrapped dropout rate as a function of pseudobulk expression (counts per million; CPM) colored by kit. **e)** Density curves of mean bootstrapped gene specificity scores colored by kit. **f)** Scatter plot of half-maximal gene drop-out (GD50) rates vs. peak gene specificity scores across 500 paired bootstrap iterations colored by kit.

Mitochondrial (MT) percentage is a feature often considered in scRNAseq analyses because it can both inform biological function ^16,17^ and serve as an indicator of cellular stress ^18,19,20,21^. Across the evaluated kits, we observed significant differences in MT percentage per cell (Kruskal-Wallis P value < 0.01) (**Fig. 2b**, left panel). In particular, BD displayed a substantially higher MT percentage compared to other kits, which is in agreement with a recently published evaluation ^22^. This may be due to unique lysis conditions that disrupt both the cell and MT membranes or lower cell viability after loading on device. Conversely, the 10x FRP, Parse, and Scale kits showed relatively low MT percentages (post-hoc Dunn’s BF-adj. P value < 0.01) **(Supplementary Tables 9-10)**. Notably, 10x FRP targets only 11 out of 13 MT genes (**Supplementary Table 11**). However, even with a comparison of MT percentage using the 11 MT genes detected in all kits, the 10x FRP kit still demonstrated reduced MT percentage. (**Extended Data Fig. 4b**). The remaining variation in MT percentage may be due to kit-specific experimental attributes such as cell processing and lysis buffer. Ribosomal protein (RP) gene percentage is another important feature because RP transcripts typically account for a large portion of sequencing reads^23^ and differential RP expression can distinguish cellular function ^23,24^. Most kits exhibited RP percentages around 10% of overall transcripts, while 10x FRP, Parse, and Scale exhibited significantly lower RP content (Kruskal-Wallis P value < 0.01; post-hoc Dunn’s BF-adj. P value < 0.01) **(Fig. 2b**, right panel; **Supplementary Tables 9, 12).** For the 10x FRP kit, this observation is consistent with the fact that it does not contain probes against RP transcripts. The relatively low RP fraction exhibited by Parse has also been reported in a separate comparative analysis ^10^.

The 10x FRP, Parse, and Scale kits are unique among the evaluated approaches because they exhibited relatively low MT and RP percentages in addition to being the only protocols that require cell fixation and permeabilization. Previous studies that investigated the effect of cell fixation and permeabilization on scRNA-seq performance show a similar reduction in RP and MT fractions for fixed cells relative to fresh cells ^25,26,27^. These studies posit that fixation and permeabilization can lead to loss of transcripts from the cytosolic compartment. Because the cytoplasm harbors the majority of both RP and MT mRNAs, a corollary of this process would be the relative over-representation of nuclear transcripts in fixed cells. Several studies have suggested that a distinguishing feature of nuclear transcripts is a relatively high degree of intron retention due to incomplete splicing ^28,29,12^. Therefore, the observation that both Parse and Scale exhibit relatively high intronic versus exonic read alignments is consistent with the expected over-representation of nuclear transcripts due to fixation and permeabilization (**Extended Data Fig. 4c).** While the evidence is incomplete in the case of 10x FRP, which lacks intron-targeting probes, these results suggest that the observed reduction in both RP and MT read fraction may also be due to the fixation and permeabilization process.

Next, we evaluated the relative gene expression across kits at the pseudobulk level. There were 11,186 genes detected with all kits, while the number of genes private to individual kits ranged from 1 (Scipio) to approximately 350 (Fluent and Scale) (**Extended Data Fig. 4d, Supplementary Table 13**). Considering only commonly detected genes, pseudobulk quantitative expression (mean UMI counts per million) profiles were calculated for each sample. We observed excellent correlation among replicates (median Spearman r = 0.99) and comparisons between kits exhibited good agreement (median Spearman r = 0.85). The combinatorial kits generally displayed lower correlations with other methods (median Spearman rho = 0.75), particularly versus Honeycomb and Scipio (Spearman r ∼ 0.62). To identify distinguishing gene expression factors, we performed principal component analysis (PCA) followed by hierarchical clustering of the first two principal component (PC) embeddings. This revealed that both the combinatorial kits and 10x FRP are located at the extremes of each axis, apart from the remaining kits (**Fig. 2c and Extended Data Fig. 4e**).

Using the PC loadings, we investigated the impact of gene length, gene GC content, and total UMI counts on expression. Strikingly, PC1 loadings exhibited a strong positive correlation with gene length (Pearson r = 0.68), which partly determines the distribution of kit embeddings along the PC1 axis (**Supplementary Table 14**). In particular, 10x FRP and the combinatorial kits are on opposite extremes of the PC1 axis, indicating that gene length can explain a portion of the differential expression between these groups. Conversely, we observed a negative correlation between PC1 and GC content (Pearson *r* = −0.41). However, there is substantial collinearity between gene length and GC content (Pearson *r* = −0.57), so the relative influence of these factors is hard to determine (**Extended Data Fig. 4f)**. Because total UMI counts impact scRNA-seq expression quantitation, we investigated the correlation between PC loadings and pseudobulk total UMI counts per kit. This revealed a strong positive correlation between total UMI counts and PC2 (Pearson r = 0.60), suggesting that library complexity has a significant effect on expression analysis.

For PC2, we noticed that protocols requiring fixation and permeabilization (10x FRP, Parse, and Scale) aggregate at the positive end of the axis, consistent with a moderate negative correlation between MT expression and PC2 embeddings (Pearson r = −0.42) and a much weaker correlation to PC1 embeddings (Pearson r = 0.02) (**Supplementary Table 14**). Gene set enrichment analysis (GSEA) using PC loadings queried against Biological Process (BP) gene sets from the Gene Ontology (GO) collection corroborates the inverse proportionality between MT content and PC2 loadings. Gene sets associated with mitochondrial functions like “proton transmembrane transport”, “electron transport chain” and “cellular respiration” were among the significant negatively enriched terms (**Extended Data Fig. 4g**). Overall, these results suggest that technical factors can have differential impacts across kits with respect to both global and pathway-specific gene expression.

Next, we investigated the stability of gene expression among the kits, specifically focusing on the drop-out rate of each gene. The drop-out rate was computed using bootstrapped cell populations, sampling from aggregated replicates of each kit. The observed exponential decay of drop-out rate relative to psuedobulk expression indicates that the proportion of non-zero expressing cells increases with higher aggregate expression level (**Fig. 2d**). Because drop-out rate only captures the binary variation between expressing versus non-expressing cells, we also computed the gene specificity score (an extension of Shannon entropy)^30^, which evaluates gene expression as a continuous variable. Thus, going beyond drop-out rate, gene specificity is able to also capture the variation in gene expression level. Using the aforementioned bootstrapped cell populations, gene specificity distributions were computed for each kit to evaluate the variability of expression within and among cells (**Fig. 2e**). We observed that pseudobulk expression at half-maximal gene drop-out rate (GD50) non-linearly and positively correlates with gene specificity distribution peaks. Notably, the bootstrapped sample sets of each kit were discretely localized (**Fig. 2f**). Hierarchical clustering of these values identified three tiers: *Tier 1)* 10x FRP, BD, 10x 3’, 10x 5’, and Parse *Tier 2)* Fluent, Honeycomb, Scale, and *Tier 3)* Scipio (**Extended Data Figure 4h**). The Tier 1 kits demonstrated the highest gene expression stability, which translates to improved variance estimates that impact downstream statistical tests, such as those used for differential gene expression analysis.

### Cell composition

To systematically annotate cell types across different kits (**Fig. 3a)**, we first labeled cells using the agreement of two independent reference-to-query label transfer methods, CellTypist ^31^ and Seurat Label Transfer ^32^, trained on an annotated PBMC dataset derived from various scRNA-seq technologies containing ten cell type labels ^12^ **(Extended Data Fig. 5a-c)**. The label agreement procedure demonstrated 94% precision, 81% recall, 86% F1 score (all weighted by class size) and an overall accuracy of 81% on a validation set (**Supplementary Table 15**). We then refined the initial cell labels using Leiden clustering and local smoothing of integrated cells across all of the kits **(Extended Data Fig. 5d-j)**. This approach leverages similar expression patterns exhibited by cells within Leiden clusters to recover unassigned cells (discordant labels between Seurat and CellTypist) and to update cell labels according to neighborhood affiliation, yielding 96.2% cell annotation across all kits (**Extended Data Fig. 5k**).

**Figure 3.**
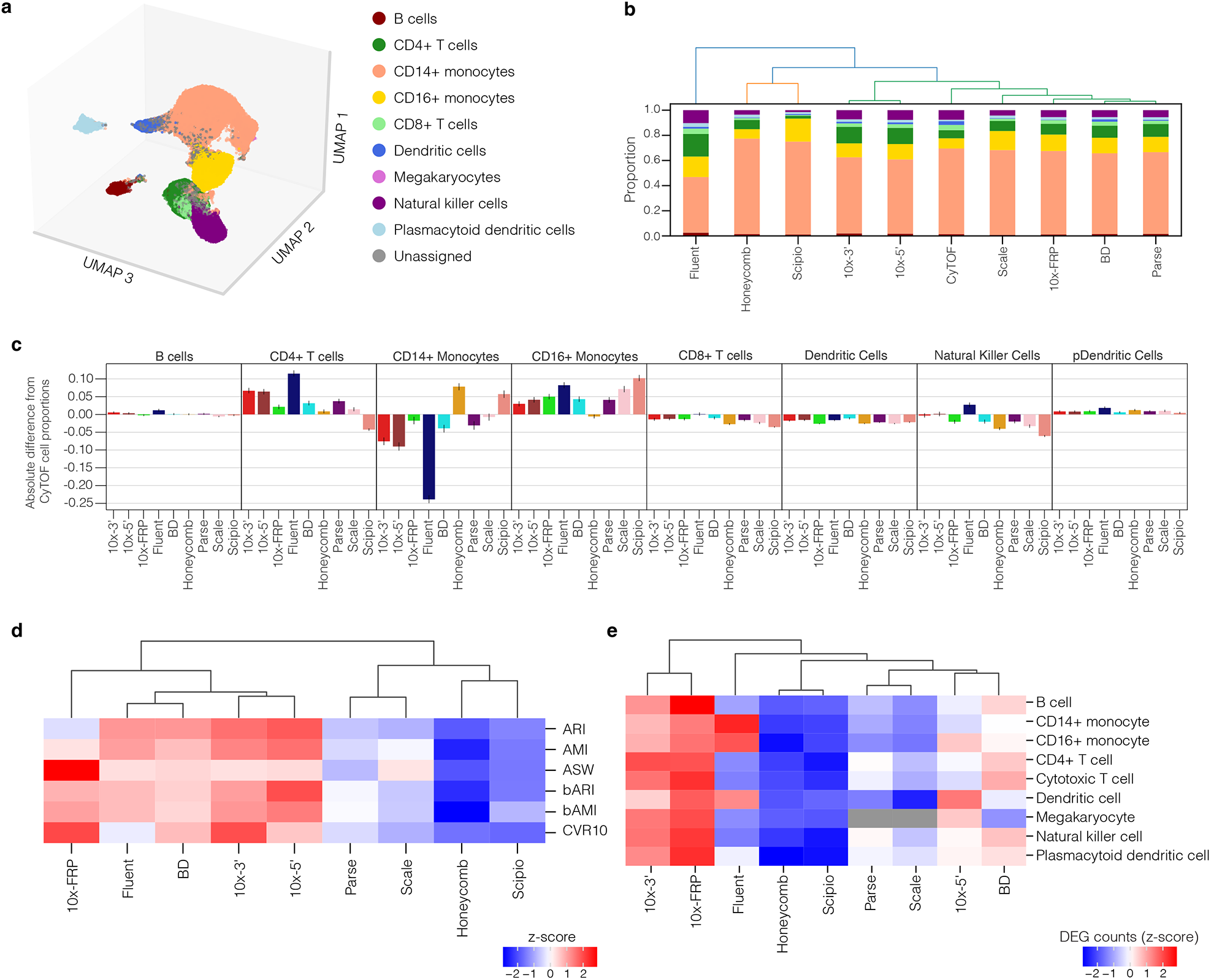
Cell type distribution, partitioning, and differential gene expression analysis. **a)** 3-dimensional uniform manifold approximation and projection (UMAP) of integrated data, colored by annotated cell type. Unassigned cells are those with discordant labels between Seurat and CellTypist that could not be resolved with Leiden cluster smoothing **b)** Mean bootstrapped cell type distribution ordered by hierarchical clustering dendrogram. Cell types are colored according to the legend in Figure 3a. **c)** Mean bootstrapped absolute difference from CyTOF reference cell proportions for each cell type and method. Error bars represent the 95% bootstrapped confidence interval. **d)** z-score of cell partitioning metrics, organized by hierarchical clustering dendrogram. Metrics included: balanced Adjusted Mutual Information (bAMI), balanced Adjusted Rand Index (bARI), Adjusted Mutual Information (AMI), Adjusted Rand Index (ARI), Average Silhouette Width (ASW), and Cumulative Variance Ratio (CVR). **e)** Mean DEG count z-scores per cell type across, ordered by hierarchical clustering dendrogram.

We aimed to assess the fidelity of cell type proportions determined by each kit relative to those obtained with CyTOF. With accuracy and specificity comparable to standard flow cytometry ^14^, CyTOF was used to ascertain the reference distribution of cell types based on 40 markers targeting PBMCs. Hierarchical clustering of bootstrapped cell type proportions derived from each kit revealed that Fluent, Honeycomb, and Scipio exhibit large deviations from the CyTOF reference distribution, especially for CD14+ monocytes (**Fig. 3b,c**). Strikingly, Fluent exhibited the largest magnitude deviations in absolute proportion relative to CyTOF: nearly 25% fewer CD14+ monocytes and greater than 10% more CD4+ T cells.

### Cluster discrimination

For each scRNA-seq kit, we sought to assess the partitioning of cell types into discrete, homogenous groups. Toward this end, data generated using each kit were individually processed to identify cell embeddings based on kit-specific transcriptomic features. Using these embeddings, we assessed various performance metrics for distinguishing cell type groups: balanced Adjusted Mutual Information (bAMI), balanced Adjusted Rand Index (bARI)^33^, Adjusted Mutual Information (AMI), Adjusted Rand Index (ARI), Average Silhouette Width (ASW), and Cumulative Variance Ratio (CVR) (see methods section for details on each metric). These metrics approach a value of 1.0 when cells segregate into well-resolved and homogenous groupings reflective of cell types. We then performed hierarchical clustering on the z-scores for these metrics to compare kits (**Fig. 3d**). These analyses revealed that 10x 3’, 10x 5’, 10x FRP, Fluent and BD exhibit relatively high scores, associated with well-resolved cell labels in discrete clusters (**Extended Data Fig. 5l,m**), in contrast to Honeycomb, Parse, Scale and Scipio **(Supplementary Table 16)**.

Next, we sought to determine the impact of kit-specific cell partitioning on differential gene expression among cell types. We reasoned that kits capable of generating discrete and homogeneous cell groupings would result in more sensitive detection of differentially expressed genes (DEGs). Across all cell types, 10x FRP and 10x 3’ yielded the highest number of DEGs, while Honeycomb and Scipio exhibited the lowest number of DEGs (**Fig. 3e, Supplementary Table 17)**. Interestingly, Fluent exhibited high variation across cell types, showing abundant DEGs for monocytes and dendritic cells but few for all other cell types.

## Discussion

Recent advancements in scRNAseq technologies have enabled researchers to choose between a diverse set of technologies to meet their specific investigative needs. However, with this diversification, choosing between available technologies can pose a challenge. Using a PBMC sample from a single donor blood draw, we performed a systematic comparison of currently available commercial kits to characterize features and performance metrics relevant to selecting an appropriate scRNA-seq approach. To facilitate the comparison of kits, we have provided a summary of relevant features in addition to library-, gene-, and cell-level performance metrics (**Table 1**, **Fig. 4**).

**Figure 4.**
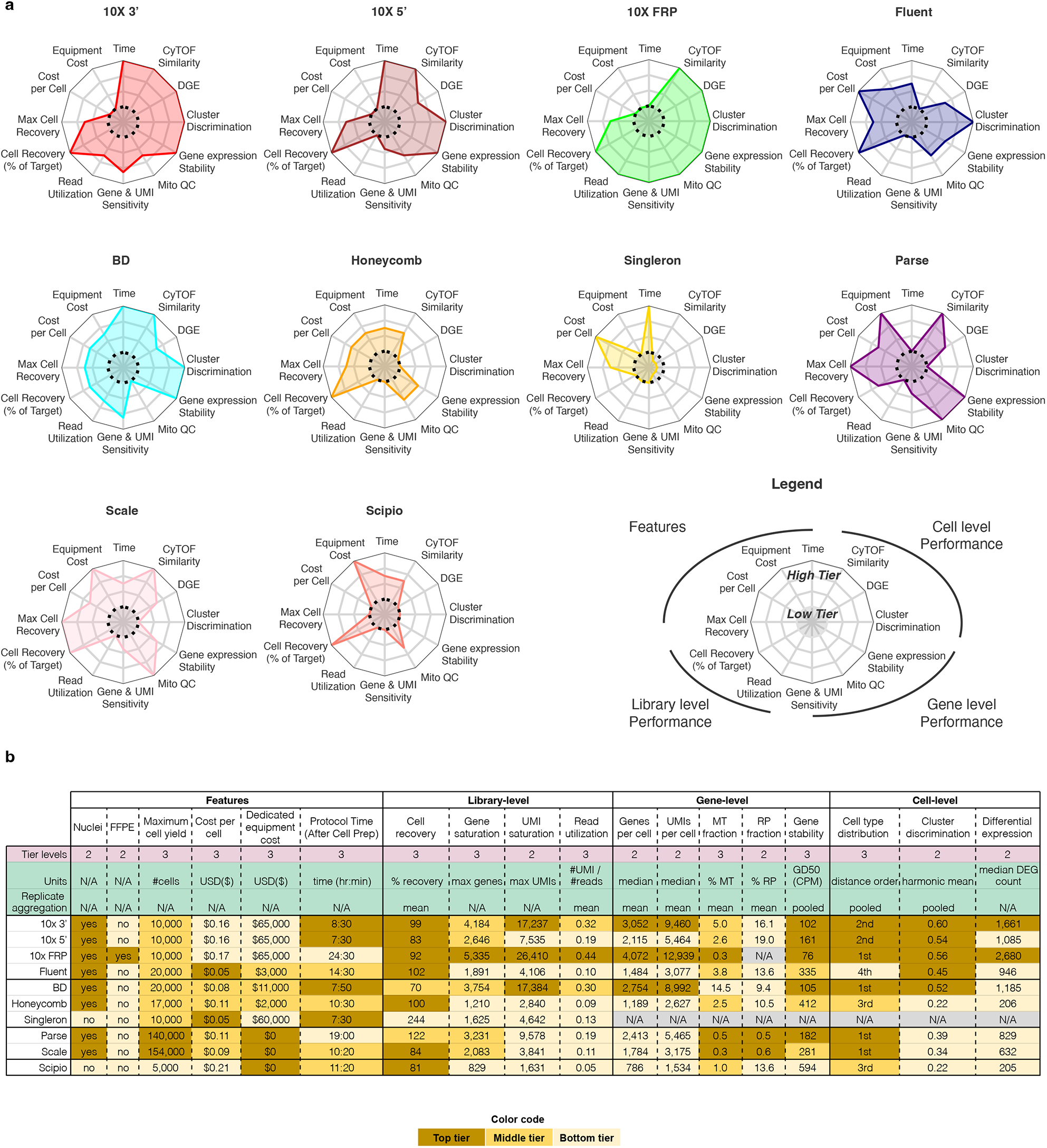
Summary of key features and performance results. **a)** Relative performance on metrics important to kit selection (rankings: inner ring = low tier, outer ring = high tier, gray region = N/A). Key metrics are grouped into four categories. **b)** Table of metric values colored by rankings and broken into groups. Details about the tiering, units, and aggregation method are provided at the top.

Overall, 10x FRP exhibited the highest ranked performance followed by 10x 3’, BD, Scale, 10x 5’ / Parse (tie), Fluent, Honeycomb, and Scipio. Singleron was excluded from the cell-based comparisons due to unreliable cell calling; however, based on UMI count, gene count, and usable mapped reads, it exhibited the lowest performance at the library level. Interestingly, at the gene level, we found a significant reduction in MT and RP fractions in kits that require cell fixation and permeabilization (10x FRP, Parse, and Scale). We hypothesized that cell fixation and permeabilization lead to loss of cytosolic transcripts including MT and RP transcripts, which was corroborated by an observed increase in the proportion of intronic reads likely derived from unspliced transcripts in the nucleus. Additionally, we noticed that BD yielded a distinguishingly high MT fraction. Because relatively high MT and RP fractions are commonly used to identify low-quality cells in scRNA-seq data, kits exhibiting lower MT and RP fractions were ranked favorably in **Figure 4**. Note that all results are based on a single PBMC sample, so the observed variation in MT and RP fractions between kits is the result of technical factors (*e.g.* missing probes, fixation, etc.); therefore, it is important to select cell quality thresholds that account for kit-specific biases. Lastly, while most kits exhibited consistent ranked performance across individual cell-level metrics, Fluent displayed high variation. Despite exhibiting the highest deviation in cell type distribution relative to CyTOF, Fluent demonstrated discrete cell partitioning and moderate differential gene expression sensitivity.

With respect to the transcript detection method, 10x FRP is unique among the evaluated kits because it utilizes probe hybridization to RNA rather than RT. While the use of probe hybridization is associated with the superior performance demonstrated by 10x FRP, it is currently limited to the detection of target coding regions in human and mouse genomes. In addition, probe hybridization is currently incompatible with characterization of expressed genetic variation, which precludes the application of genotype-based demultiplexing tools ^34,35^. Lastly, the variable number of probes and target sequences for each gene may introduce bias in expression quantitation. For RT-based kits, the extent of gene body read coverage is critical to genotyping and specialized applications such as TCR/BCR repertoire analysis. While most RT-based kits exhibited biased coverage to either the 3’- or 5’-end of genes, Parse and Honeycomb demonstrated relatively uniform coverage throughout gene bodies due to the use of random hexamers during cDNA generation (**Extended Data Fig. 6**).

Another set of distinguishing features relates to sample type compatibility. Here, again, 10x FRP is distinct due to its ability to process cells, nuclei, and formalin-fixed paraffin-embedded (FFPE) tissue. Its unique ability to process FFPE tissue greatly expands the reach of scRNA-seq to access the vast catalogs of preserved clinical specimens. Relatedly, 10x FRP and the combinatorial indexing kits, Parse and Scale, require sample fixation, which enables asynchronous sample acquisition and storage followed by batch processing. In general, batch processing of either fixed or fresh samples improves efficiency and increases scale. For example, a high-throughput variant of the 10x FRP kit can be configured with barcoded probe sets to enable simultaneous processing of up to 16 samples while still allowing the capture of up to 10,000 cells per sample. Similarly for Parse and Scale, plate-based sample barcoding and combinatorial indexing enables batch processing of up to 96 samples. However, the labor-intensive process of split-pool barcoding requires a high number of input cells due to sample loss during successive rounds of washing and barcoding. Another multiplexing technique is cell hashing which utilizes barcoded antibodies originally developed for emulsion-based methods^36^ such as 10x 3’ and 10x 5’ but is amenable to other scRNA-seq technology groups. In addition, cell hashing enables the detection of multiplets, which is especially useful for maximizing cell recovery yield^19^ and further reducing cost per cell.

While analytical performance and features define assay utility, cost is among the most salient factors in selecting a scRNA-seq kit. To compare kits using a common basis, we computed cost per cell by dividing kit cost by the claimed maximum cell yield, excluding the cost of sequencing (**Fig. 4b)**. This analysis revealed Fluent and Singleron as the most cost-efficient kits, followed by BD and Scale, Honeycomb and Parse, the 10x variants, and lastly Scipio. It is worth noting that higher-throughput kit versions along with specific experimental configurations, such as cell overloading, can further reduce cost by increasing cell recovery yield. Additionally, there are upfront equipment costs associated with each kit that contribute to expense assessment and these costs vary drastically among kits. For instance, the dedicated equipment for Singleron and the 10x kits range around $60,000-65,000 while Scipio, Parse, and Scale do not require any specialized equipment (**Table 1)**. Therefore, a holistic expense assessment must consider the costs of both procuring equipment in addition to projected kit consumption over time. When averaging upfront equipment and kit costs over expected output ranging from 10^4^-10^7^ cells, the Fluent kit emerges as the most cost-efficient; whereas, the 10x Genomics platform expenses generally exceed all other kits (except Scipio beyond about 1.3 million cells) (**Supplementary Table 18a**). Importantly, these cost estimates are based on the features of the currently evaluated kits. The use of higher throughput variants of these kits (*e.g.* 16-ple× 10x FRP) together with other multiplexing strategies will dramatically impact the cost structure. Moreover, even though we omitted sequencing costs in our calculations, we identified substantial variation in aspects of read utilization among kits, which can impact cost per cell. For instance, about 90% of reads from the 10x FRP kit were usable (mapped and CB-tagged); whereas, Fluent utilizes less than 40% of reads. This inefficiency essentially doubles the sequencing cost per cell to achieve the same number of usable reads.

We estimated protocol duration based on published user guides, starting from either fresh or fixed cells and ending with completed libraries, which revealed that current commercially-available scRNA-seq kits require at least about 8 hours (10x 3’, 10x 5’, BD, and Singleron) and as long as 24.5 hours (10x FRP; accounting for an overnight incubation step). Analysis of the cell processing rate (maximum cell yield over protocol duration) indicates that the 10x FRP protocol requires about 36-fold more time than Scale, the most time-efficient kit, to process an equivalent number of cells (**Supplementary Table 18b**). Depending upon anticipated experimental throughput, factors such as upfront equipment cost, kit cost, and protocol time are important variables in selecting an appropriate scRNA-seq method.

This study provides a comprehensive and systematic evaluation of commercially-available scRNA-seq kits and sheds light on the various strengths and weaknesses of each kit with respect to time, cost, functionality and performance, which should facilitate an optimal and informed selection of a suitable scRNA-seq approach. However, there are important limitations to the scope of work and interpretation of results. First, while insights derived from PBMCs provide a useful perspective, other sample types such as nuclei, fixed tissue, and whole blood may reveal an alternative assessment of relative kit performance. Second, we observed an unusually high fraction of monocytes in our PBMC sample compared to the typical distribution found in healthy donors^12^; however, this over-representation of monocytes was corroborated by CyTOF and was relatively consistent across kits. Third, in order to focus resources on evaluating a broad variety of kits, we did not pursue multiplet rate analysis, which would have required additional “barnyard” experiments using mixed human and mouse cells. While we do not provide empirical multiplet rates, simulation-based estimates are provided **(Supplementary Table 19)**. Lastly, we observed substantial cell recovery variation using the pipelines provided by Fluent and Singleron, which may impact performance assessments by failing to exclude low-quality cells and ambient RNA. Given the ever-expanding set of diverse scRNA-seq methods, further research into technology-agnostic cell calling algorithms would be an important contribution to improving accuracy and consistency in the field.

In this study, we benchmarked commercially-available scRNA-seq kits to enable researchers to select the appropriate technology for their experimental needs. In addition, by identifying the strengths and weaknesses of each methodology, we expect that this knowledge will help advance scRNA-seq technology development. Beyond the current set of analyses, this work provides a rich dataset consisting of 218,154 called cells across 10 kits with over 120,000 cell type annotations. Such a dataset can be used to train annotation models or to benchmark algorithms that explicitly account for technical variability among scRNA-seq approaches. Taken together, we have established a comprehensive experimental and analytical framework for comparing scRNA-seq technologies. The insights and underlying data generated from this work have the potential to facilitate substantial progress in the field of single-cell biology.

---

**Extended Data Figure 1. Schematics of cDNA generation approaches. a)** Emulsion-based protocols. **b)** Microwell-based protocols. **c)** Combinatorial-indexing protocols. **d)** Matrigel-based protocol.

**Extended Data Figure 2. Downsampling sensitivity analysis. a)** Coefficient of variation for number of cells detected across all downsampled depths (30,000 to 2,000 in steps of 2,000 reads per cell). **b)** Number of cells recovered at each sampling depth (according to cell calling algorithms accompanying each kit). **c)** Number of cells returned at each sampling depth for Singleron when using a kernel density estimator (natural log UMI count) instead of the internal Singleron cell calling algorithm. **d)** Hierarchical clustering dendrogram for the gene saturation fitted curve. **e)** Hierarchical clustering dendrogram for the UMI saturation fitted curve. Hierarchical clustering performed on 34 equidistant fitted values along the curve of each kit.

**Extended Data Figure 3. UMI recovery and read utilization a)** Average UMI recovery in cells (total UMIs / total input FASTQ) to identify natural breaks. **b)** Average UMI recovery in cells. **c)** Components of read usage per cell: average number of input reads per cell (left), average number of mapped reads per cell (middle), and average number of recovered UMIs per cell (right).

**Extended Data Figure 4. Gene composition. a)** Hierarchical clustering dendrogram for median gene and UMI counts of the filtered and processed 30K read depth subsample data. **b)** Fraction of MT gene counts with respect to genes common to all kits. **c)** Fraction of reads mapping to exonic (left) and intronic (right) regions. Reads must align completely within either an exon or intron. **d)** Spearman correlations of pseudobulk gene expression profiles. **e)** Hierarchical clustering dendrogram of PC1 and PC2 embeddings. **f)** Collinearity analysis of gene length and gene GC-content. **g)** Gene set enrichment analysis of PC1 and PC2 loadings. **h)** Hierarchical clustering dendrogram of for mean GD50 and mean peak gene specificity scores across 500 bootstrapped samples.

**Extended Data Figure 5. Cell annotation. a)** Schematic of two-pass cell annotation strategy. **b-c)** 3D and 2D UMAPs of cells colored by first-pass cell annotation results. **d-e)** 3D and 2D UMAPs of cells colored by kit after *Harmony* integration showing mixing of kits across cell types. **f)** Integrated cell population divided into lymphocyte-like cells in blue (left, group B) and monocyte-like cells in red (right, group A). Two-pass cell annotation results for the lymphocyte-like group (middle) and monocyte-like group (right). **g-h)** For lymphocyte-like population, Leiden cluster cell type composition after first- and second-pass annotation. **i-j)** As above, for the monocyte-like population. **k)** Fraction of unassigned cells remaining after annotation (per kit). **l)** 2D PCAs of each kit (independently processed and visualized) colored by the two-pass cell annotation results. **m)** 2D UMAPs of each kit (independently processed and visualized) colored by the two-pass cell annotation results.

**Extended Data Figure 6. Metagene coverage.** Density of reads found in cells mapping to exonic regions across gene bodies. Coverage plots are organized according to technology groups: **a)** Emulsion-based protocols, **b)** Microwell-based protocols, **c)** Combinatorial indexing protocols, and **d)** Matrigel-based protocols.

## Data availability

Sequencing data were deposited to NCBI Sequence Read Archive (SRA) under the BioProject accession PRJNA1106903.

## Code availability

The code used to analyze data from this study has been deposited in the GitHub repository: https://github.com/danledinh/scRNAseq-repo

## Acknowledgements

We thank Isabel Adams for administrative assistance. In addition, we thank David Garfield, Bo Li, and Russell Xie for providing thoughtful comments.

## Contributions

M.D.S. and S.D. conceived the study. M.D.S., J.L., H.B., B.W., C.C., and H.M. performed the scRNA-seq experiments and prepared sequencing libraries and Y.L, J.M.L, M.S, A.X.M sequenced the libraries. A.A-Y., S.L. and W.E.O performed the CyTOF experiment. J.H., S.V., A.K. and D.L. performed data processing and analysis. Z.M., D.L. and S.D. supervised the project. M.D.S., J.H., D.L. and S.D. prepared the manuscript. All authors discussed the results and approved the manuscript.

## Corresponding authors

Correspondence to Daniel Le and Spyros Darmanis.

## Ethics declarations

All authors are employees of Genentech, a member of the Roche Group.

## Methods

### Sample preparation, viability assessment, and sequencing

Multiple aliquots of frozen Human Peripheral Blood Mononuclear Cells were purchased from Stem Cell Technologies (Cat No 70025.2). PMBCs were thawed in complete RPMI 1640 Medium (Thermo Fisher Scientific Cat No 11875093), washed twice in PBS, and resuspended in PBS 0.04% BSA (Thermo FIsher Scientific Cat No AM2616). Cell concentration and viability were assessed using Cellometer K2 (Nexcelom Bioscience) using dual-fluorescent staining (AO-PI). The viability of all the PBMC aliquots was above 95%. For each kit, PBMC aliquots were processed according to the manufacturer user guide. Barcoded cDNA libraries were sequenced with Illumina sequencers (NextSeq2000 and NovaSeq6000). The sequencing specifications are listed in **Table 1** in the *Sequencing Format* column.

### PBMC phenotypic characterization

The PBMC sample was analyzed using CyTOF with a panel of 37 antibodies (**Supplementary Table 20a**) ^37^. PBMC subpopulation frequencies, relative to the number of live cells, were calculated according to the gating strategy in **Supplementary Table 20b**.

### Experimental design

For the majority of evaluated kits, two replicates targeting 10,000 cells were generated using PBMC aliquots derived from a single healthy donor (Only one sample was generated for both Parse and Scale. Fluent had three replicates). Note that Scipio does not allow processing of more than 5,000 cells. PBMC aliquots were thawed directly prior to use. All assays were performed according to kit instructions. Sequencing was performed with a target depth of slightly more than 30,000 reads per cell. Downsampling of FASTQ files was needed to yield the targeted 30,000 reads per cell. All downsampling was performed by the software *seqtk* version 1.4 (https://github.com/lh3/seqtk) with the flags `sample −2 -s100`.

### Data processing pipelines

Data processing software supplied by each kit vendor was implemented to produce UMI count matrices. For each of the 10x kits, *CellRanger* version 7.1.0 was used. The 10x 3’ and 5’ kits required the subcommand `count` and 10x FRP required the subcommand `multi`. For Fluent, *PIPseeker* version 2.1 was used. For BD, *Rhapsody* version 2.0 was used. For Honeycomb, *BeeNet* version 1.1.3 was used. For Parse, *split-pipe* version 1.1.1 was used. For Scale, *ScaleRna* version 1.3.3 was used. For Singleron, *CeleSCOPE* version 1.17.0 was used with the subcommand `multi_rna`. For Scipio, *Cytonaut* version 7 was used. The human reference (Ensembl 98/GENCODE v32) from 10x Genomics (refdata-gex-GRCh38-2020-A) was used for alignment and feature quantification. This reference was adapted for compatibility with all other kit pipelines.

### Cell calling algorithms

Unlike the cell calling algorithms provided with most kits, the softwares for Fluent and Honeycomb require user input to identify cells. For the Fluent pipeline, a user can select from five outputs with different cell count yields. The outputs are based on five sensitivity levels according to the formula:

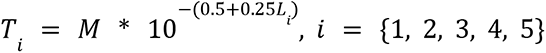

where *T* is the UMI threshold for a barcode to be called a cell, *M* is the number of UMIs for the first ranked barcode where the transcript count does not drop by more than 10% of the previous barcode, and *L* is the multiplier for each of the five sensitivities (*i = {1, 2, 3, 4, 5}*). We selected the sensitivity level that yielded cell counts closest to the targeted 10,000 cells. Honeycomb requires users to input the expected cell number and returns exactly this number of barcodes.

### Read utilization analysis

FASTQ reads were classified into three categories: A) uniquely mapped and tagged with a cell barcode, B) uniquely mapped and tagged with a non-cell barcode, and C) all other reads removed or masked by data processing pipelines (*e.g.* unmapped reads, untagged reads, duplicates). To determine the number of reads in categories A and B, the BAM files outputted from each pipeline were processed with the following command: `samtools view -F 256 “$bam_file” | grep “$bc_tag” | sed “$sed_pattern” | sort | uniq -c | sort -nr | awk ‘{print $2 “,” $1}’; $bam_file:` path to BAM file, `$bc_tag:` cell barcode tag (*e.g.* `CB:Z:`), `$sed_pattern:` pattern to extract full barcode (*e.g.* `‘s/.*CB:Z:\([ACGT]*\).*/\1/’`). The resultant set of uniquely mapped and barcoded reads were further segregated based on possession of a passing cell barcode (category A) versus a non-cell barcode (category B). The number of reads in category C was determined by subtracting A and B from the number of reads in the input FASTQ file.

### Sensitivity as a function of sequencing depth analysis

Downsampled sub-libraries were generated with total reads equivalent to the target reads per cell, ranging from 30,000 to 2,000 reads per cell at steps of 2,000. This resulted in 14 sub-libraries per sample, which were processed using respective software accompanying each kit. The output included median number of genes and UMIs per cell at each sampling depth. For each kit, aggregating across replicates, we fit a Michaelis-Menten saturation function (a specific application of the rectangular hyperbolic function) to these data ^38^. The Michaelis-Menten function is:

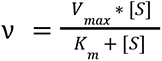

Here, ν is the median gene or UMI count per cell, *V_max_* is the saturation level, [*S*] is the read depth, and *K_m_* is the depth that yields half-maximal saturation. Curve fitting was performed using the `curve_fit()`function from the python package *scipy.optimize* version 1.13.0. Kits were grouped into sensitivity performance tiers based on hierarchical clustering of 34 equidistant expected values along fitted curves derived from each kit.

### Counts matrix preprocessing

Starting with the counts matrix derived from the 30,000 reads per cell library, cells with fewer than 5 genes were removed^39^. Doublets were removed using *Scrublet*^40^ *version 0.2.3* with a doublet prediction score threshold of 0.25 for all samples. Next, the lowest number of cells yielded by a kit targeting 10,000 cells (after minimum gene number and doublet filters; n = 7,750) was used to randomly subsample all count matrices to a uniform sample size. Scipio was excluded because its maximum yield is 5,000 cells. Lastly, genes present in fewer than 3 cells per sample were removed^39^.

### Quality control metrics

Commonly assessed metrics such as gene counts, UMI counts, mitochondrial percentage, and ribosomal protein percentage were identified on count-matched subsamples of cells (n = 7750, excluding Scipio; Scipio-rep-1: n=3457, Scipio-rep2: n=4425) passing gene and cell number controls (see *Counts matrix preprocessing*). The *scanpy* package (version 1.9.1) function `sc.pp.calculate_qc_metrics()` with flags `qc_vars=[’mt’,’ribo’]` was used to calculate counts and gene metrics. Mitochondrial and ribosomal genes have gene symbols starting with `“MT-”` or `“RPS“/“RPL“`, respectively.

### Statistical analysis of kit variables

For per-cell variables such as gene counts, UMI counts, mitochondrial percentage, and ribosomal percentage, non-parametric tests were performed to determine distributional differences among kits. We used the Kruskal-Wallis test (*scipy package*; `stats.kruskal()`) followed by post-hoc Dunn’s tests for pairwise comparisons (*scikit_posthocs* package; `posthoc_dunn()`). P values were adjusted for multiple testing using the Bonferroni procedure.

### Informative feature selection

Highly deviant genes (HDGs) were identified according to this tutorial: https://www.sc-best-practices.org/preprocessing_visualization/feature_selection.html (*scry* package; https://rdrr.io/bioc/scry/src/R/featureSelection.R). HDG determination is based on counts; therefore, they are unaffected by the inconsistent application of normalization procedures^41^. To calculate deviance, a python implementation of the deviance calculation was used (see Code availability section).

### Pseudobulk gene expression

Pseudobulk gene expression was computed from UMI counts per million (CPM) across all cells in a given sample. Specifically, pseudobulk expression was defined as mean log-transformed (`np.log1p()`) CPM. The intersection of expressed genes across all samples was used as the feature set for PCA. The resultant PC loadings (*i.e.* weights for each gene) were used for GSEA. From the *gseapy* package version 1.1.2, the function `prerank(min_size=50, max_size=1000, permutation_num=1000, seed=6)` was used to query PC loadings against the MSigDB C5 collection, GOBP (Gene Ontology, Biological Process) subset: https://www.gsea-msigdb.org/gsea/msigdb/download_file.jsp?filePath=/msigdb/release/2023.2.Hs/c5.go.bp.v2023.2.Hs.symbols.gmt. For correlations between PC embeddings and gene set expression scores, the *Scanpy* function `tl.score_genes()` was used to compute scores for each pseudobulk sample.

### Dropout rate and gene specificity scores

Drop-out rates and gene specificity scores^30^ were computed for each kit using a bootstrap subsample analysis on all cells aggregated by kit (N_boot=500, N_sample=7,750). Only the intersection of genes detected in all kits was considered. For each bootstrap iteration, subsampled count matrices were CPM normalized prior to gene drop-out and gene specificity computation. Each gene specificity bootstrap iteration was paired to the corresponding drop-out iteration to allow for subsequent comparisons.

For the gene drop-out bootstrap, we calculated drop-out rate as the fraction of cells with CPM values of zero. A drop-out rate of zero indicates that no cells express the gene while a rate of one indicates that all cells express the gene. The drop-out rate was modeled as a function of pseudobulk gene expression (mean expression per gene across all cells; CPM) using the following exponential decay equation^9^:

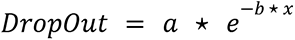

Where *x* is the pseudobulk gene expression (CPM) for a given gene, *b* is the decay rate parameter, and *a* is the scale parameter that defines the max drop-out value. The scale parameter *a* was set to a value of 1 to reflect complete dropout at zero pseudobulk expression. Curve fitting was performed in each bootstrap iteration using the `curve_fit()`function from the python package *scipy.optimize* version 1.13.0. To optimize curve fitting, initial value estimate for *b* was randomly initialized to a value between 0 and 0.05. The *b* parameter was bounded between 0 and 0.05. Estimated parameters across all bootstraps were used to compute bootstrapped mean estimates and 95% confidence intervals. Half-maximal gene drop-out values (GD50) for each bootstrap iteration were computed as follows:

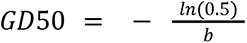

For the gene specificity bootstrap, specificity was computed according to the methodology by Martinez and Reyes-Valdes^30^ using a python implementation of the `entropySpecificity()` function in the *BioQC R* package^42^. Gene specificity for a given gene *i* is an extension of Shannon entropy and is defined as:

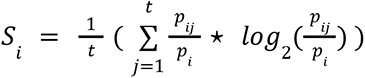

Where *S_i_* is the gene specificity score for a given gene, *t* is the number of cells in a sample (7,750 for this bootstrap), *p_ij_* is the proportion of CPM from gene *i* in cell *j*. The following equation defines the average CPM proportion of gene *i* across all cells, *p_i_* :

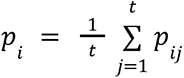

For each kit, gene specificity score densities across all bootstrap iterations were plotted using `sns.kdeplot()` from *seaborn* and `stats.gaussian_kde()` from *scipy* was used to find the distribution peak.

### Integration of cells across kits

The *Harmony* (*harmonypy version 0.0.9*) integration algorithm was used to generate adjusted PCA embeddings that account for kit-specific variation^41,43^. The top 2,000 batch-corrected HDGs across all samples were selected as the feature set. Specifically, deviance was computed per sample (batch) and the median deviance across all samples was used to rank genes. Next, normalization and scaling were performed independently for each sample (batch-specific processing) using the following *Scanpy* version 1.9.1 functions with default settings: 1) `pp.normalize_total(),` 2) `pp.log1p()`, and 3) `pp.scale()`. The integration step was performed using the *Scanpy* wrapper function (*scanpy.external*) `pp.harmony_integrate()`; `key = “$kit_id“` (*i.e. Anndata* object `.obs` column name for kit identities), `basis = “X_pca“`, `adjusted_basis = “X_pca_harmony“`). The resulting adjusted PCA embeddings were used to construct a neighborhood graph for UMAP visualization of all integrated cells: `sc.pp.neighbors(); n_neighbors=100, n_pcs=15`.

### Cell annotation

The *SeuratData R* package version 0.22 was used to access a dataset of annotated PBMCs derived from several scRNA-seq methods^12^ (loaded using `data(“pbmcsca”)`; data version: 3.0.0). This reference set consists of ten cell type labels: B cells, CD14+ monocytes, CD16+ monocytes, CD4+ T cells, CD8+ T cells (called Cytotoxic T cells in the dataset), Natural killer cells, Plasmacytoid dendritic cells, Dendritic cells, Megakaryocytes, and Unassigned cells. The reference cells were split into training and validation sets. The validation set was created by random subsampling to yield 7,750 cells to match the number of cells in each kit. The training set consisted of 20,840 cells. These data sets were used to train reference-to-query label transfer models.

Cells were annotated using the consensus of two independent label transfer methods, CellTypist ^31^ and Seurat’s Label Transfer ^32^. Cells with discordant labels across the two methods were labeled “Unassigned”. Next, the cells from all kits were integrated using the *Harmony* algorithm. The cells segregated into two coarse groups consisting of mostly lymphocytes and monocytes. For each coarse group, Leiden overclustering (*i.e.* clusters >> labels; n∼20) was performed to yield high-granularity subpopulations primarily composed of a single dominant cell type label. The majority label within each subcluster was used to relabel all constituent cells. The following provides detailed methodology:

### *CellTypist* version 1.6.2

The implementation of *CellTypist* was based on this tutorial: https://colab.research.google.com/github/Teichlab/celltypist/blob/main/docs/notebook/celltypist_tutorial_cv.ipynb. Each query (*i.e.* an individual sample dataset) was processed independently as follows: 1) 2,000 HDGs from the query were identified, 2) UMI counts per cell were normalized to 10,000 (*scanpy* package; `pp.normalize_per_cell(); counts_per_cell_after=10000),` and 3) counts were log1p transformed. Scaling was performed by *CellTypist*. Batch-normalization (*i.e.* across scRNA-seq methods) of the reference set was performed similarly to above. During the *CellTypist* model training on the reference set, approximately 1,300 (*i.e.* intersection of top 300 explanatory variables for each label) of the original 2,000 HDGs were selected as the final feature set. Label transfer we performed using the argument `majority_vote = True`.

### *Seurat Label Transfer* version 4.1.1 (*R* version 4.2.0)

Label transfer was based on this *Seurat* tutorial:

https://satijalab.org/seurat/articles/integration_mapping.html. The reference dataset was processed as follows: 1) data was partitioned by method, 2) each reference method was independently normalized (`NormalizeData`; default settings), 3) and the top 2,000 variable genes were identified (`FindVariableFeatures`; `selection.method = “vst”, nfeatures = 2000`). The first 30 principal components were used to find anchors for reference data integration (`FindIntegrationAnchors`; `dims = 1:30, scale = TRUE, reduction = “cca”`). For each query dataset, raw counts were normalized and the first 30 principal components were used for label transfer (`FindTransferAnchors`; `dims = 1:30, reference.reduction = “pca“` and `TransferData`; `dims = 1:30`). *SeuratDisk* version 0.0.0.9020 was used to interconvert between *Seurat* and *Scanpy* objects.

### Leiden overclustering and reclassification

After consensus annotation, an overclustering approach we implemented similar to the majority voting method employed by *CellTypist*: 1) The integrated cell population was divided into two coarse-grain groups composed of mostly lymphocytes and monocytes (`sc.tl.leiden(); res = 0.002`), 2) Each group was independently reprocessed to yield normalized and scaled data, and 3) For each of these groups, the Leiden algorithm was used to overcluster cells into at least 20 clusters. To determine the resolution needed for at least 20 clusters, a 30,000 cell subsample was used to more efficiently scan across a range of resolutions. The initial resolution was 1.25 and incremented in steps of 0.05. The resolution values of 1.65 and 1.35 were found to be optimal resolutions for the lymphocyte and monocyte subsamples, respectively. These resolutions were used to initiate a scan on the full dataset, which found resolutions 1.8 and 1.4 to yield at least 20 clusters for the full lymphocyte and monocyte groups, respectively. All cells within a fine-grained cluster were subject to reclassification if two conditions were satisfied: 1) the majority (at least 60%) of cells shared the same consensus label and 2) the majority cell type was not “Unassigned”. Otherwise, the original consensus labels remained unchanged.

### Cell type distribution analysis

For each kit, bootstrapped cell type proportions were calculated by resampling (with replacement) aggregated replicates. Resampling was performed 500 times to yield subsamples consisting of 7,882 cells (equivalent to the number of cells found in the sparsest multiple replicate dataset). Mean cell type proportions were calculated using the 500 subsamples. With respect to mean cell type proportions, hierarchical clustering (*scipy* package; `cluster.hierarchy.linkage(); method=’ward’`) was used to compare kits in relation to CyTOF. Megakaryocytes and Unassigned cells were excluded since these labels were not included in the CyTOF annotation. In addition, using the 500 subsamples for each kit, mean absolute difference relative to CyTOF cell type proportions were calculated.

### Cell partitioning scores

Kit replicates were concatenated using respective filtered count matrices (*anndata* package; `ad.concat()`). For each kit, the intersection of deviant genes across replicates were used as features in PCA (*scanpy* package; `tl.pca(); svd_solver=’arpack’, use_highly_variable=True`). Concatenated datasets were integrated using Harmony (*scanpy external* package ; `pp.harmony_integrate(); max_iter_harmony = 30`). Then, each dataset was subsampled to the number of cells found in the sparsest multiple replicate dataset (*scanpy* package; `pp.subsample()`, n = 7,882). This was followed by construction of a neighborhood graph (*scanpy* package; `pp.neighbors(); n_neighbors=30, n_pcs=15`). Leiden clustering was performed over a series of resolutions ranging from 0.02 to 1.00 in increments of 0.02 (*scib* package; `metrics.cluster_optimal_resolution()`). This process identified the optimum lowest resolution that yielded the smallest difference between the expected number of clusters and labels. All subsequent grouping-based scores were derived from clustering results at the optimum resolution of each kit. Each metric was performed using the following set of tools: ARI (*sklearn* package`; adjusted_rand_score()`), AMI (*sklearn* package`; adjusted_mutual_info_score()`), ASW (*scib* package`; metrics.silhouette()`), bARI (*balanced_clustering* package`; balanced_adjusted_rand_index()`), bAMI (*balanced_clustering* package`; balanced_adjusted_mutual_info()`), and CVR was computed as the sum of the first 10 PCA variance ratios.

### Differential gene expression (DGE) analysis

Using the filtered count matrix with cell type annotations, each sample was processed as follows: 1) The number of cells was subsampled to n=3,457 (*i.e.* the lowest cell yield; Scipio replicate 1), 2) Any cell type with fewer than 10 cells was removed from subsequent DEG analysis, and 3) A Wilcoxon rank-sum test was performed for each query cell type versus the remaining cell types (*scanpy* package`; tl.rank_genes_groups(); method = “wilcoxon“`). Annotation of significant differential expression required log2 fold-change > 2 and Benjimini-Hochberg-adjusted P value < 0.05. For DEG counts aggregated by kit rather than individual samples, mean DEG counts were provided. For each cell type, Z-score normalization was performed using the DEG counts distribution across kits (*scipy* package; `stats.zscore()`). Hierarchical clustering of kits with respect to cell-type specific DEG count Z-scores was performed using the *seaborn* package (`clustermap(); standard_scale = None`)

### Tiering system and radar plot visualization

For cell recovery tiers, observed yields within 20% of the target number of cells were classified as *Tier 1*. All other kits, excluding Singerlon, are in *Tier 2*. Singleron was classified as *Tier 3* because of cell calling discrepancies. For cost per cell, equipment cost, and read utilization a kernel density estimator (*scipy* package; `stats.gaussian_kde()`) was used to identify natural breaks (*scipy package;* `signal.find_peaks()`). For equipment cost, a natural log transform was applied (log1p). For maximum cell recovery the data were divided into three tiers: *Tier 1)* >20,000 cells, B) 10,000 - 20,000 cells, and *Tier 3)* <10,000 cells. For protocol time, the data were divided into three tiers: *Tier 1)* approximately 1 work day (< 8.5 hours), *Tier 2)* approximately 2 work days (8.5-16 hours), and *Tier 3)* more than 2 days (>16 hours). For mitochondrial fraction, the data were divided into three tiers: *Tier 1)* Approximately 0%, *Tier 2)* <10%, and *Tier 3)* >10%. For ribosomal protein fraction, the data were divided into two tiers: *Tier 1)* Approximately 0% and *Tier 2) >*0% (most were around 10%). For the remaining performance categories, hierarchical clustering was used to identify performance groupings. For genes and UMIs per cell, hierarchical clustering was performed using both median genes and UMIs per cell.

Radar plots were used to visualize the performance tiers of several categories: Protocol Time, Cost (Equipment), Cost (per Cell), Max Cell Recovery, Cell Recovery (% of Target), Read Utilization, Gene & UMI sensitivity (Gene Rank + UMI Rank), Mito QC, Cluster Discrimination, Differential Gene Expression (DGE), and Cytof Similarity (Cell Type Distribution). The performance tiers were converted to normalized scores as follows:

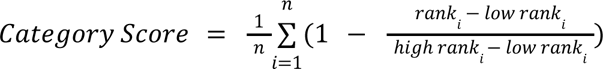

Where *n* is the number of kits, *i* indexes the kits, *rank_i_* is the tiering rank of a given kit, *low rank_i_* is the lowest numerical tier value (corresponding to best performance), and *high rank_i_* is the highest numerical tier value (corresponding worst performance). The output of this scoring system ranges from 0 (worst) to 1 (best).

### Metagene read coverage

Using *pysam* version 0.22.0, CB-tagged BAM files derived from each data processing pipeline were filtered for reads containing barcodes from valid cells. The resultant filtered BAM files were used to generate read coverage BigWig (.bw) files via the subcommand `bamCoverage` from *deeptools* version 3.5.0. Next, the .bw files were processed using subcommands `computeMatrix (--skipZeros --metagene)` and `plotProfile` to visualize average read coverage profiles across genes.

### Use of Large Language Models (LLMs)

An enterprise implementation of Chat-GPT was used for proofreading sections of the manuscript and for generating data visualization code. All LLM outputs were reviewed for accuracy.

