## Extended Data Figures for "Comparative Analysis of Commercial Single-Cell RNA Sequencing Technologies"

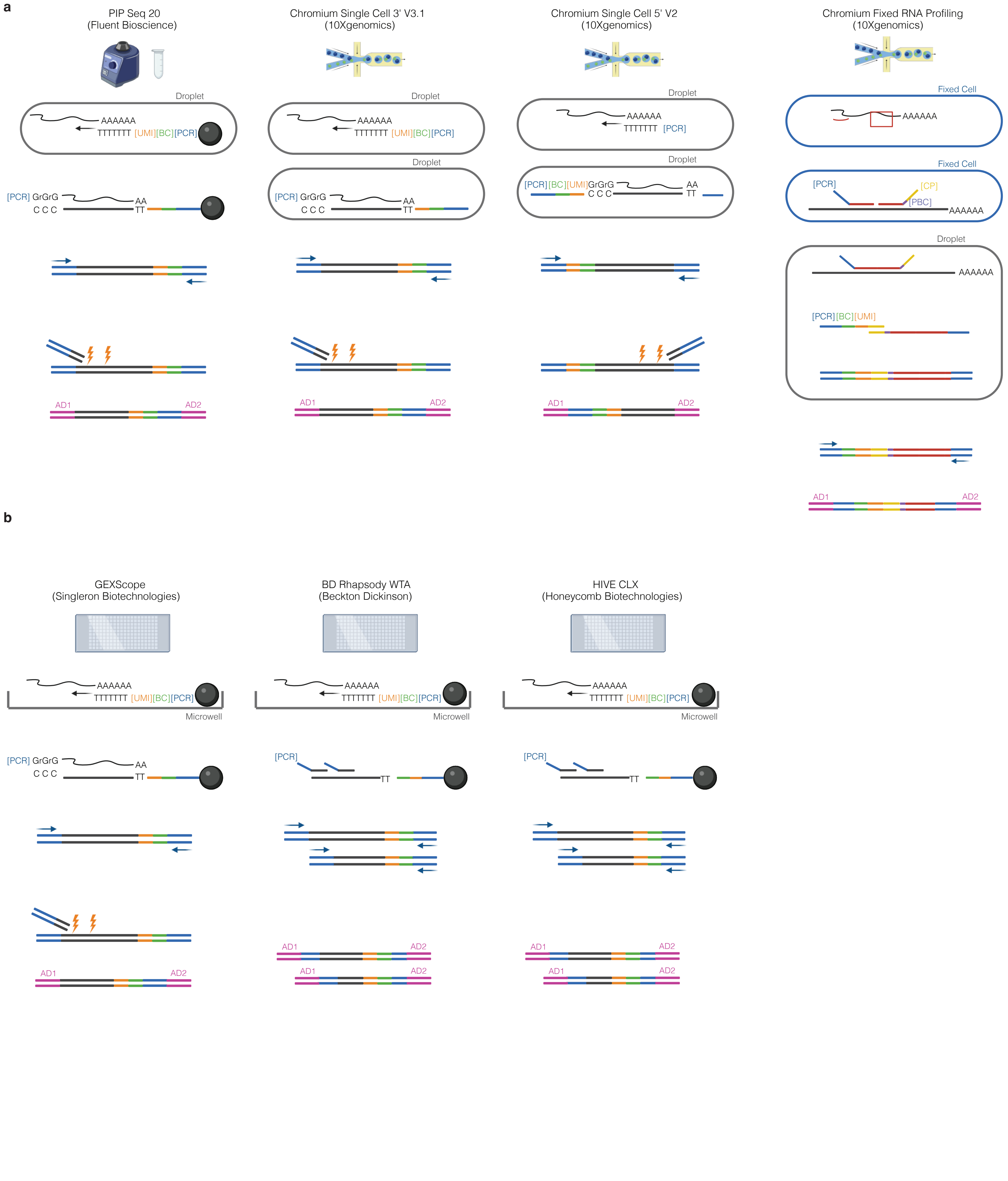


Extended Data Figure 1

c

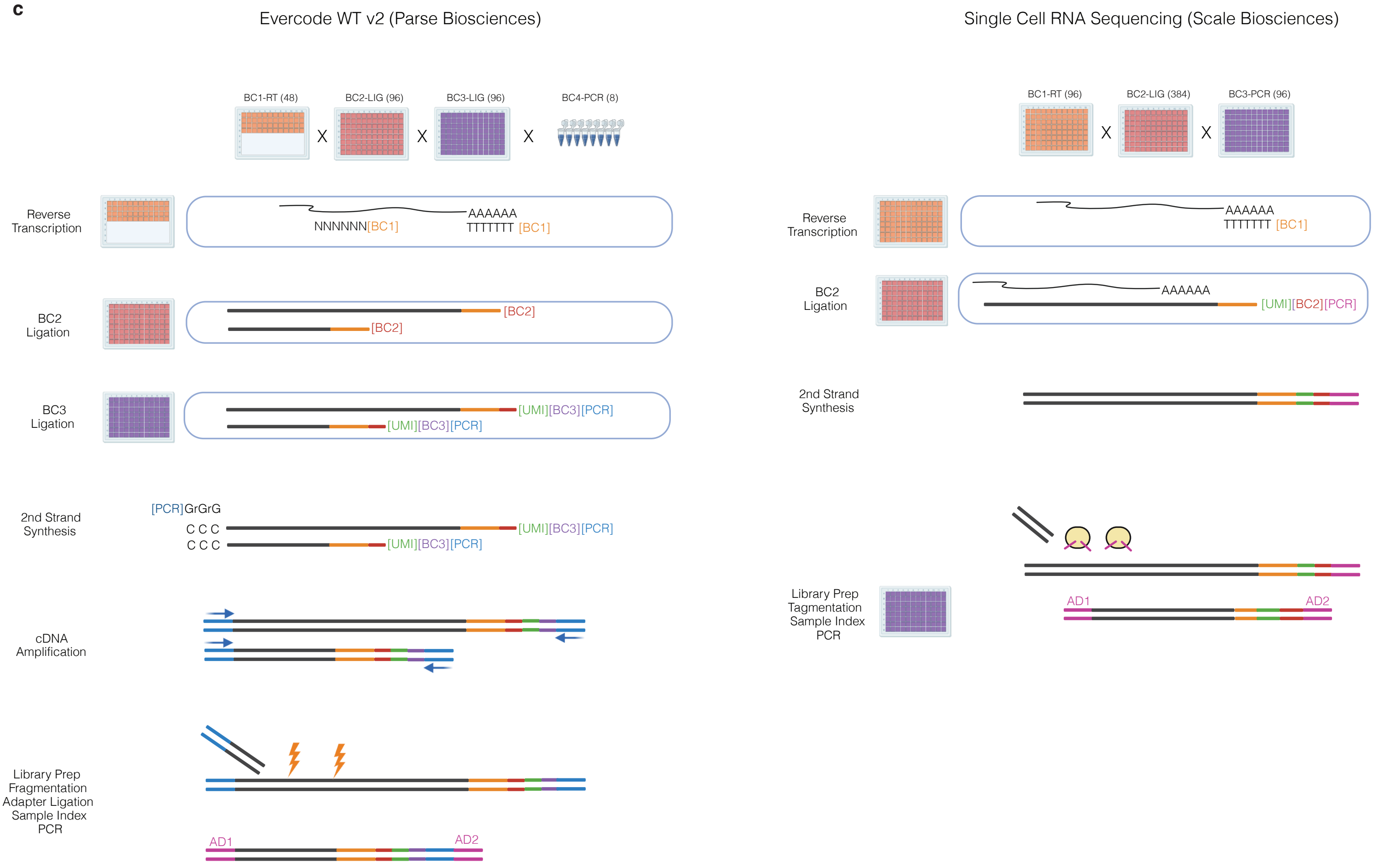


d

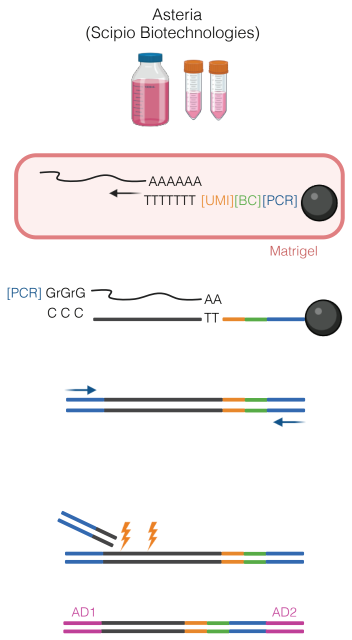


**Extended Data Figure 1**

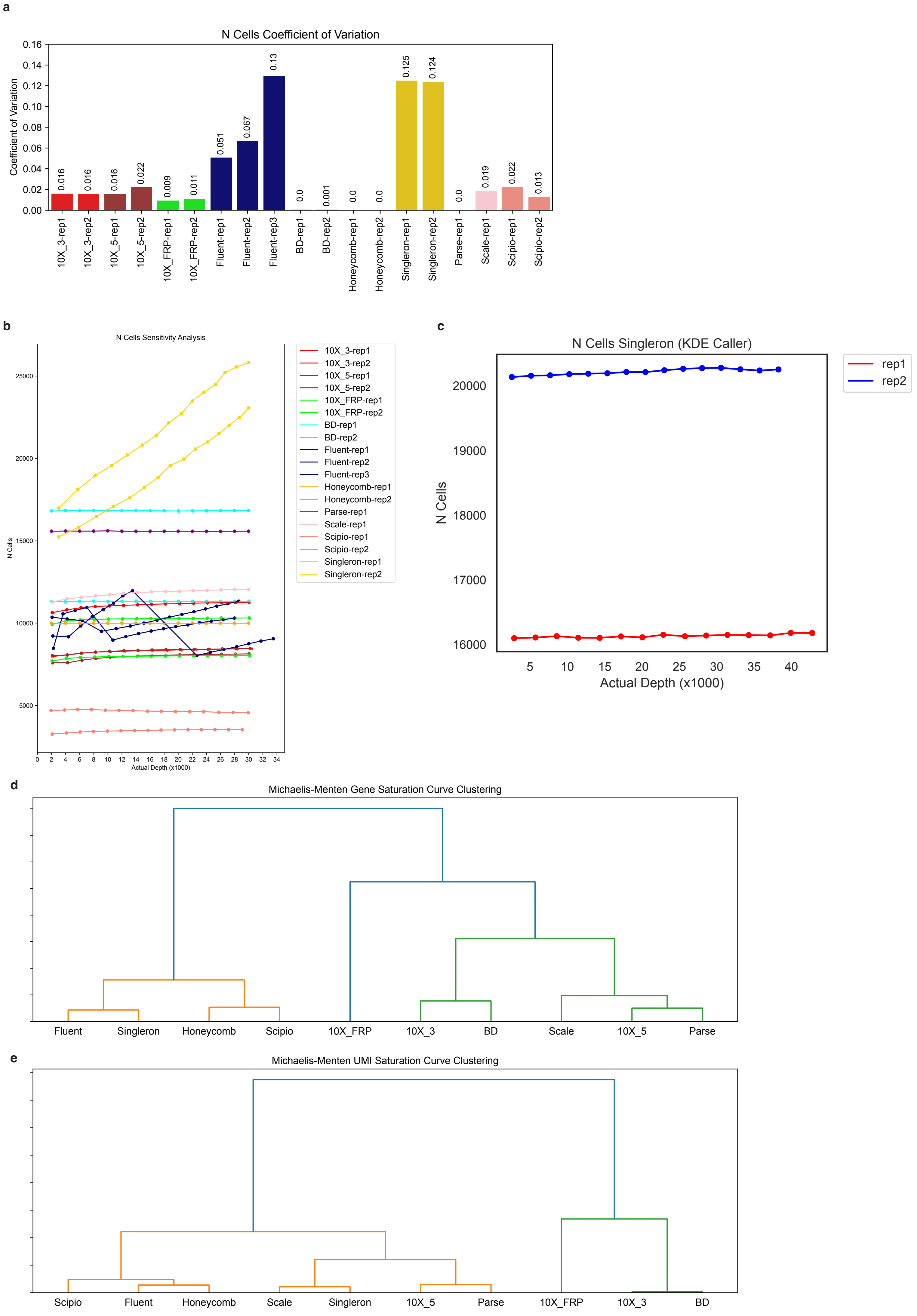


**Extended Data Figure 2**

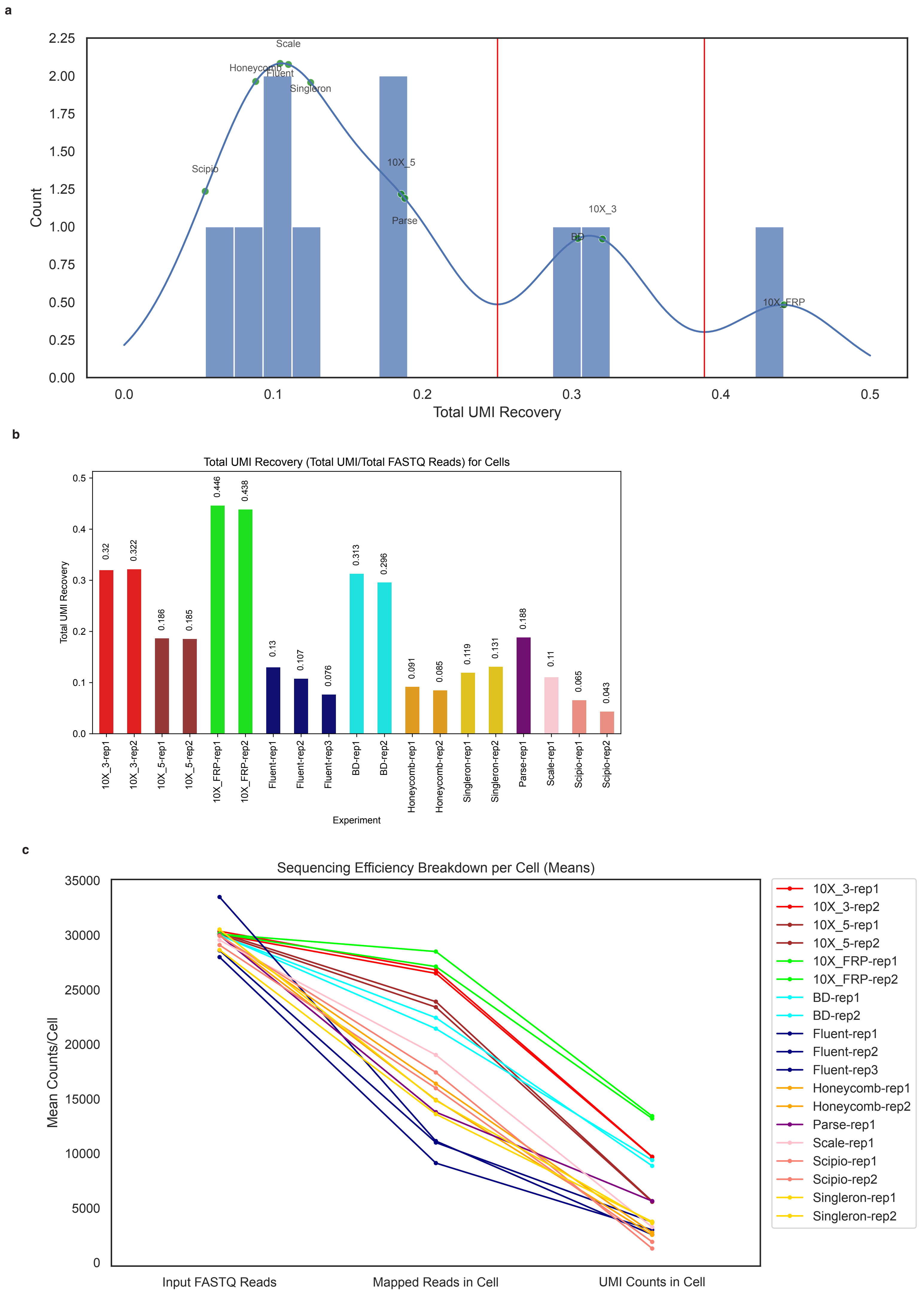


**Extended Data Figure 3**

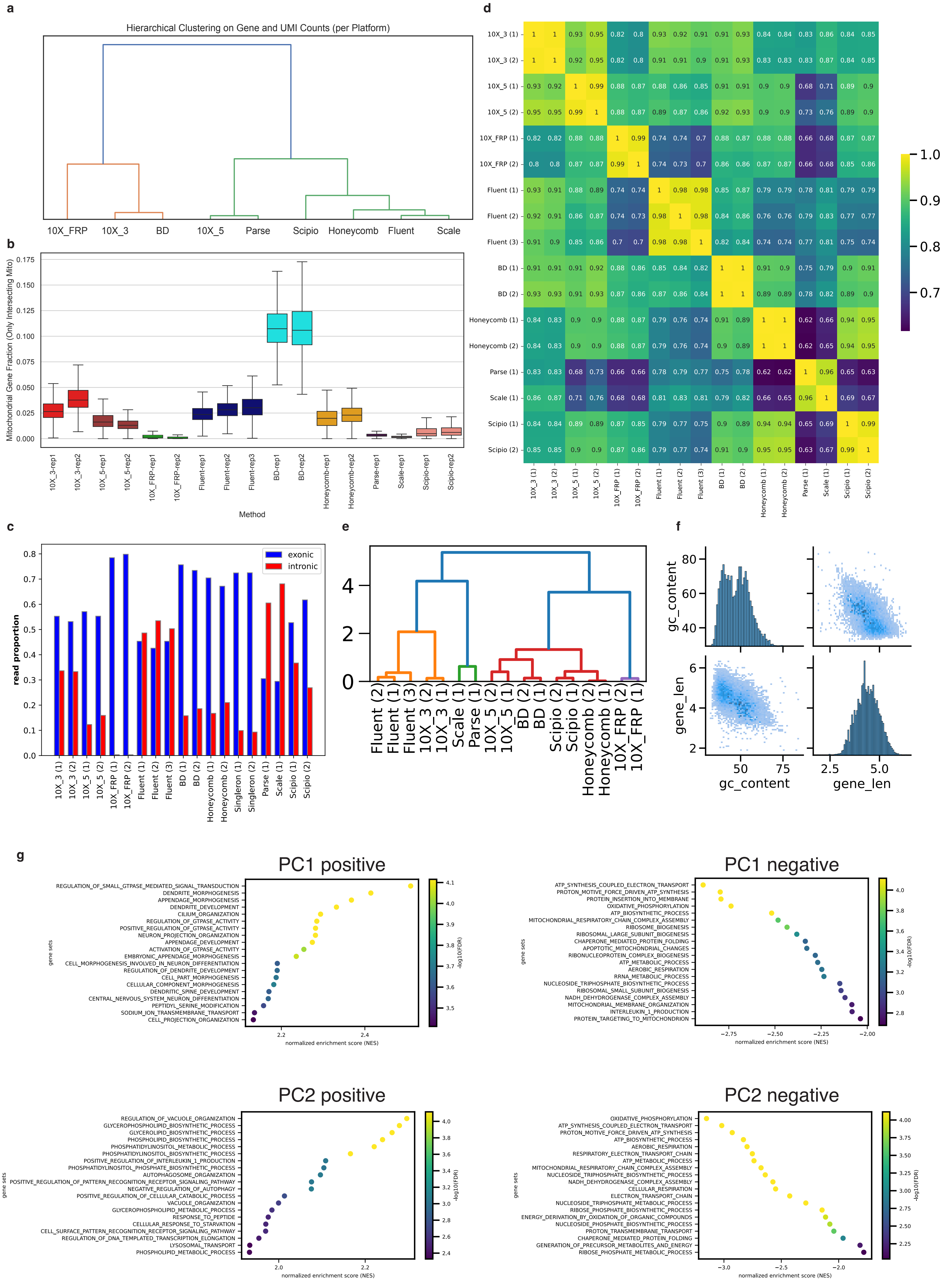


**Extended Data Figure 4**

**h**

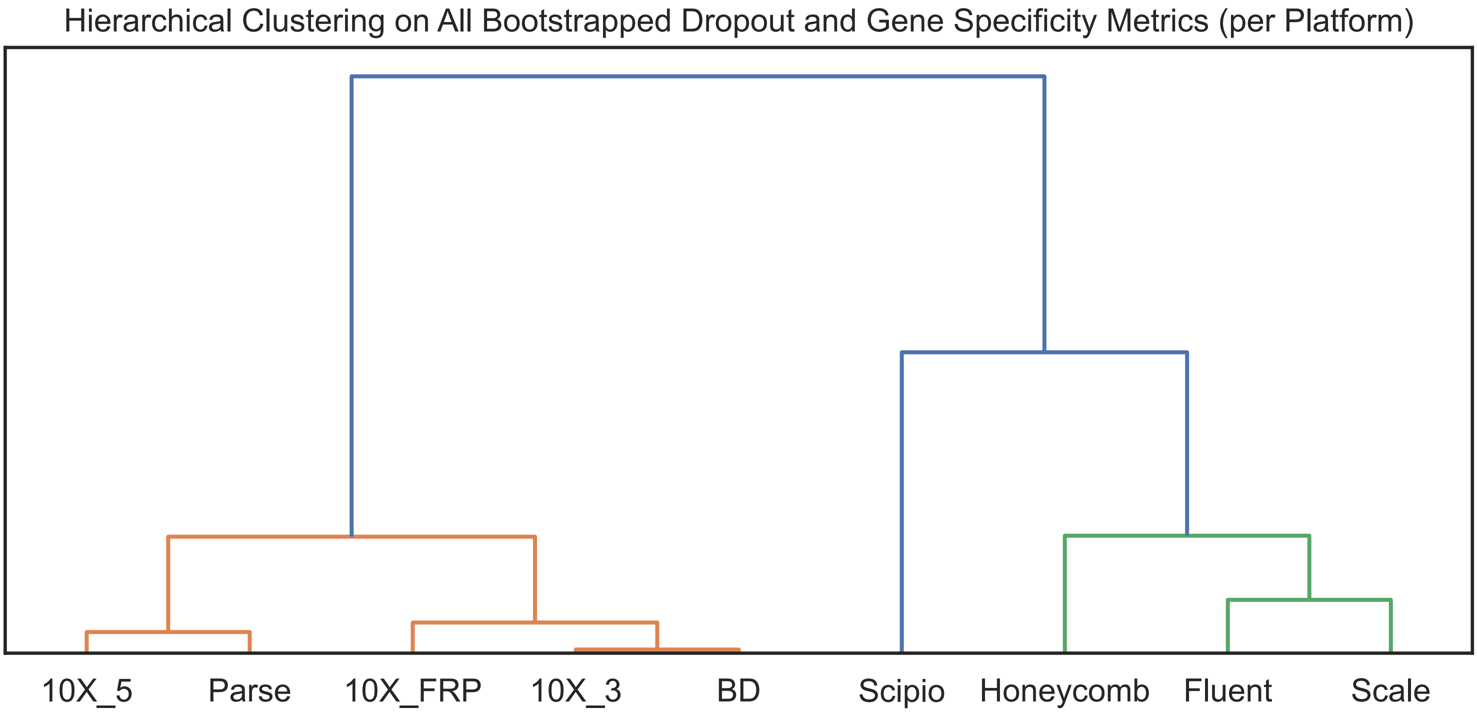


**Extended Data Figure 4**

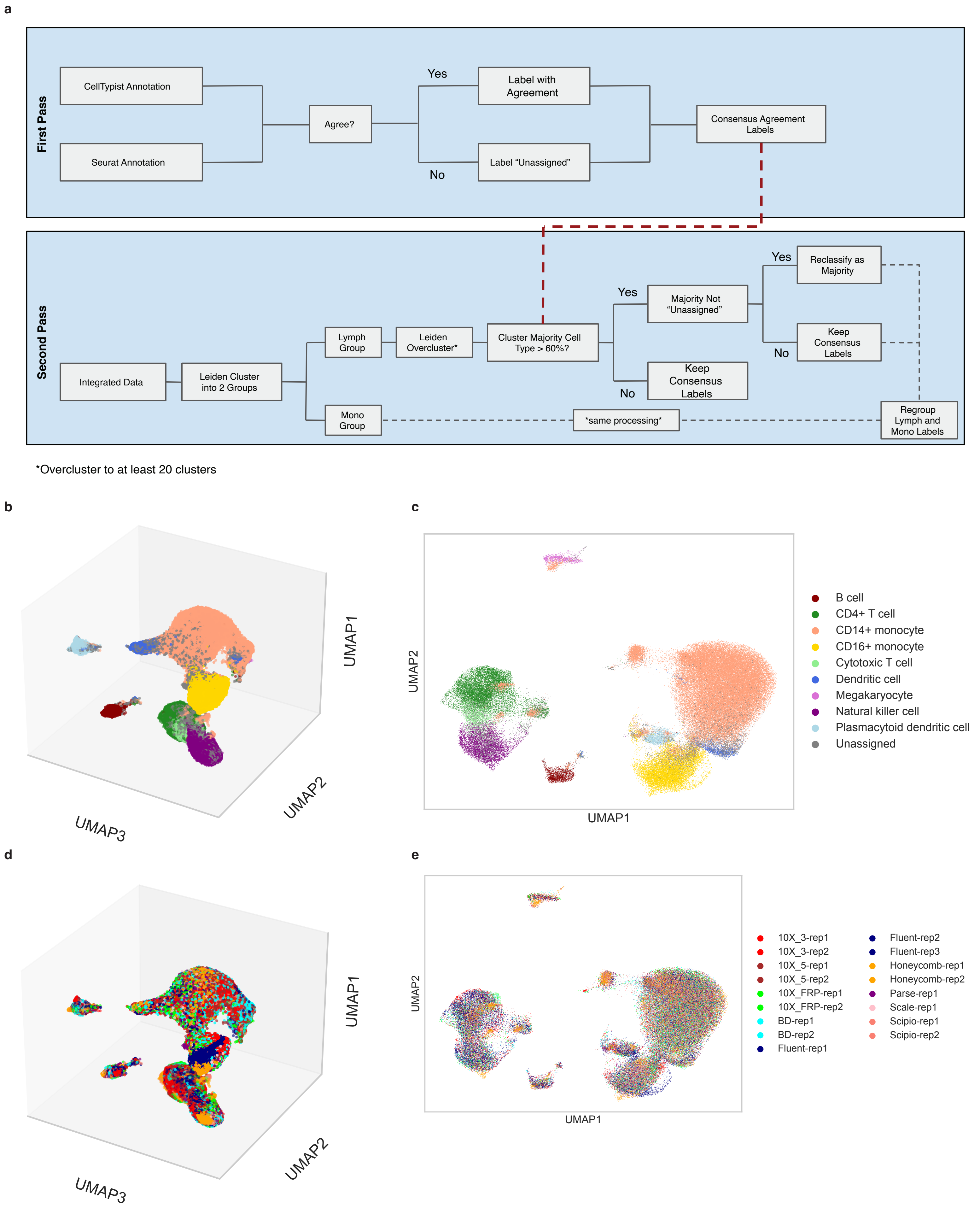


**Extended Data Figure 5**

f

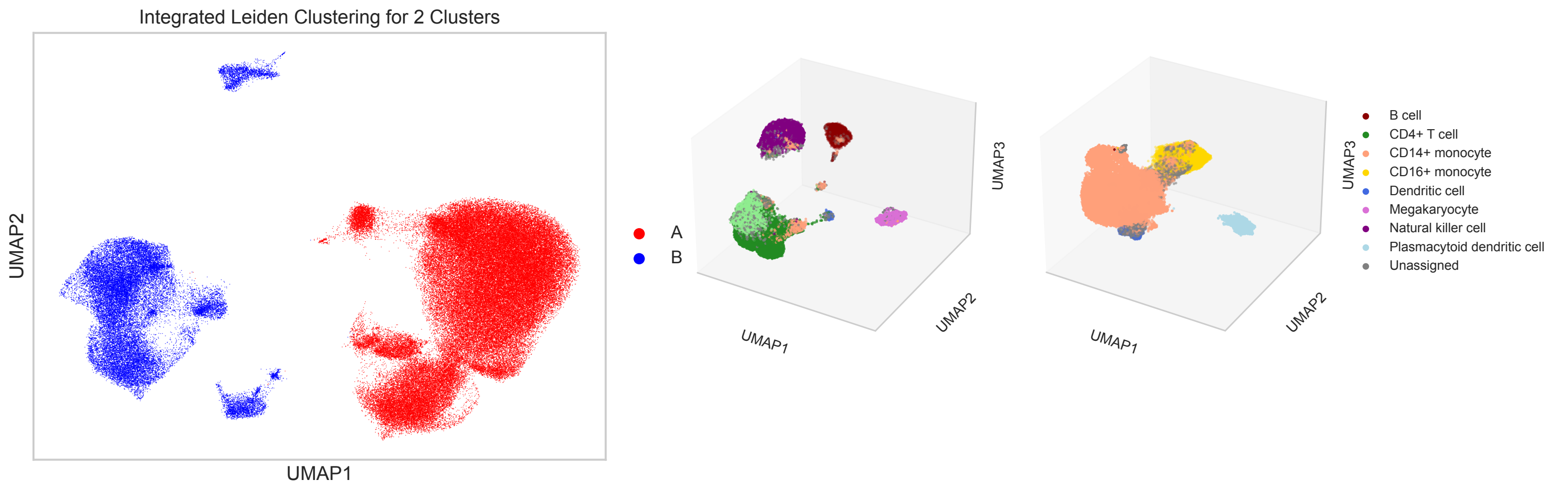


g

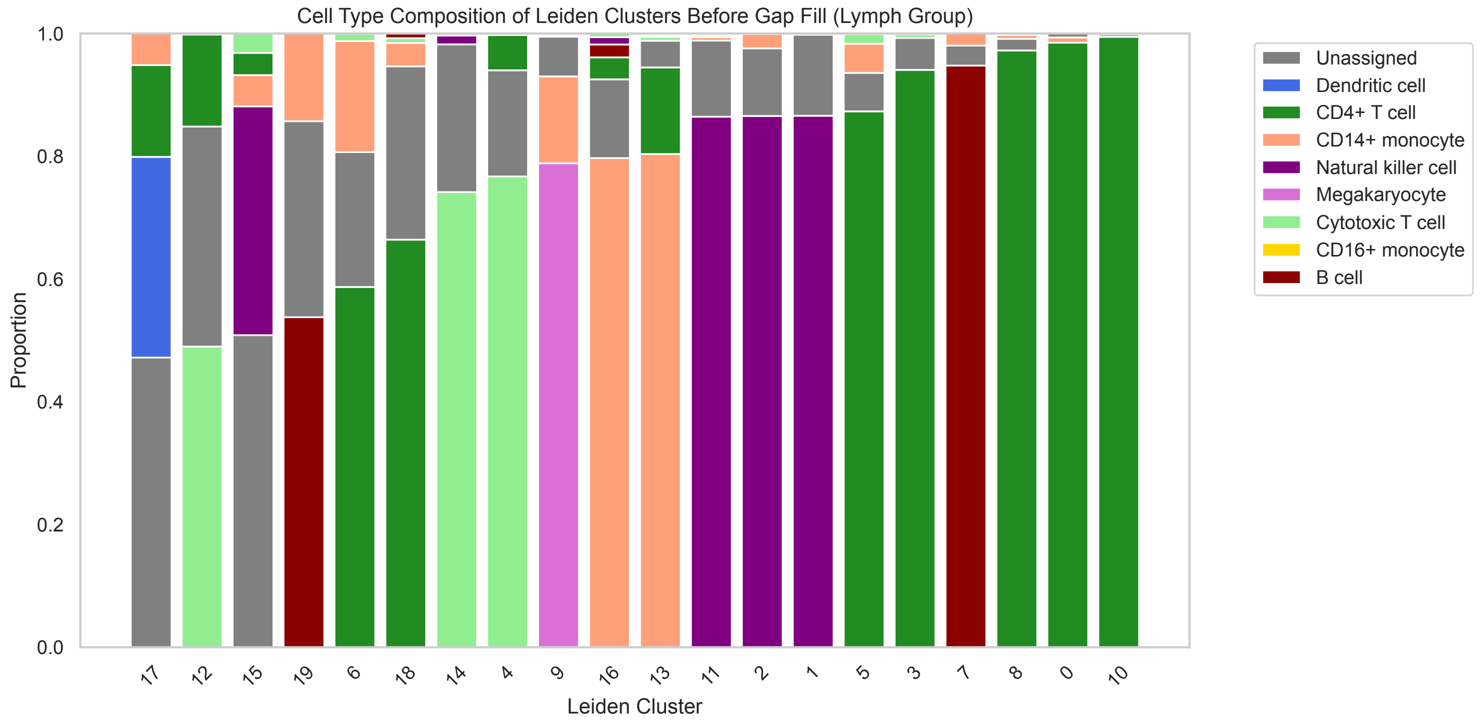


h

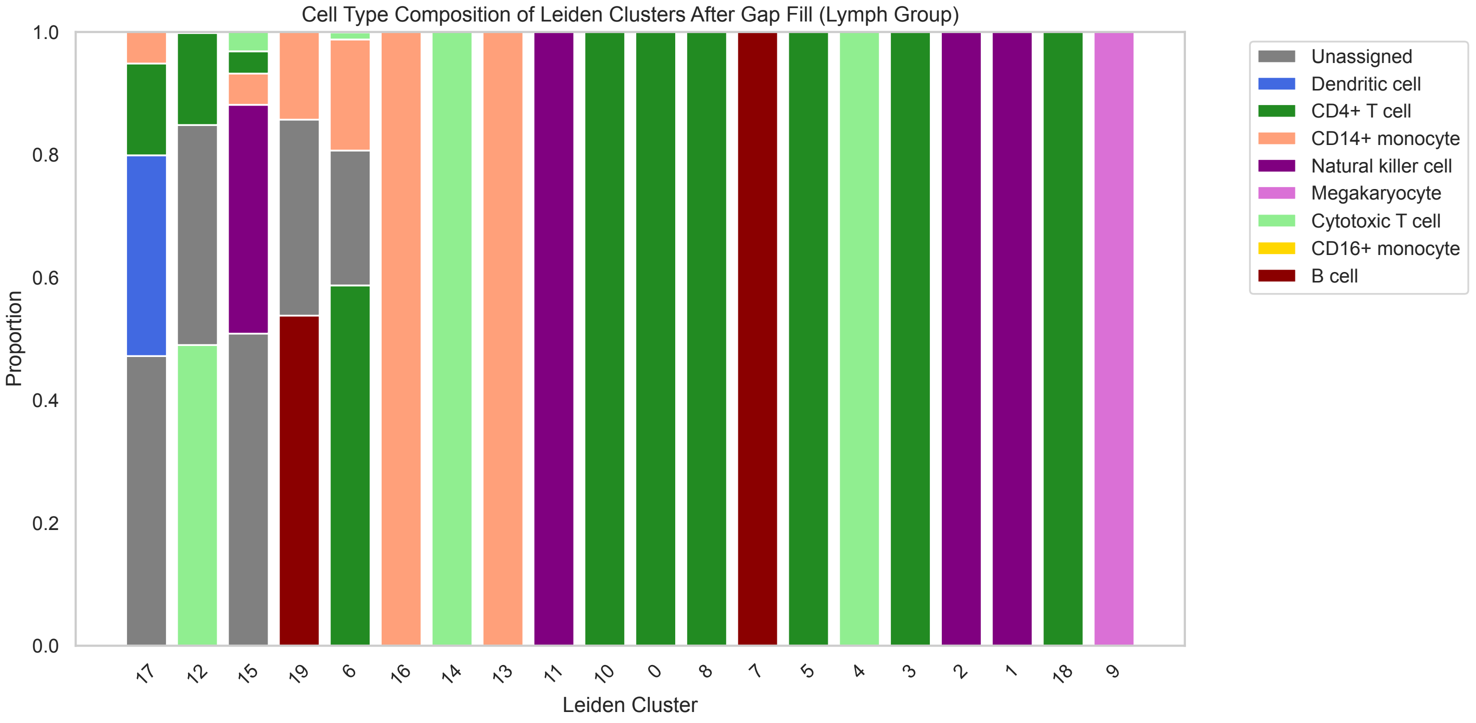


**Extended Data Figure 5**

**i**

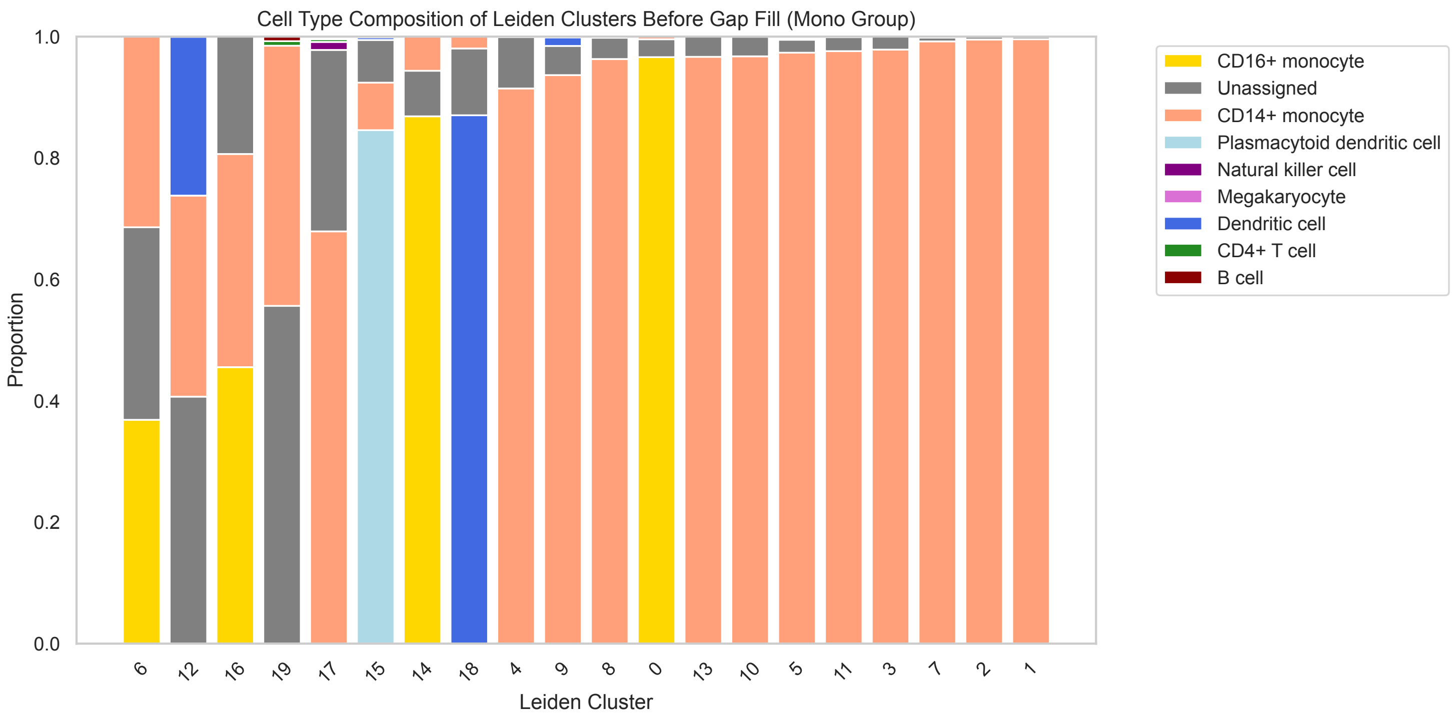


**j**

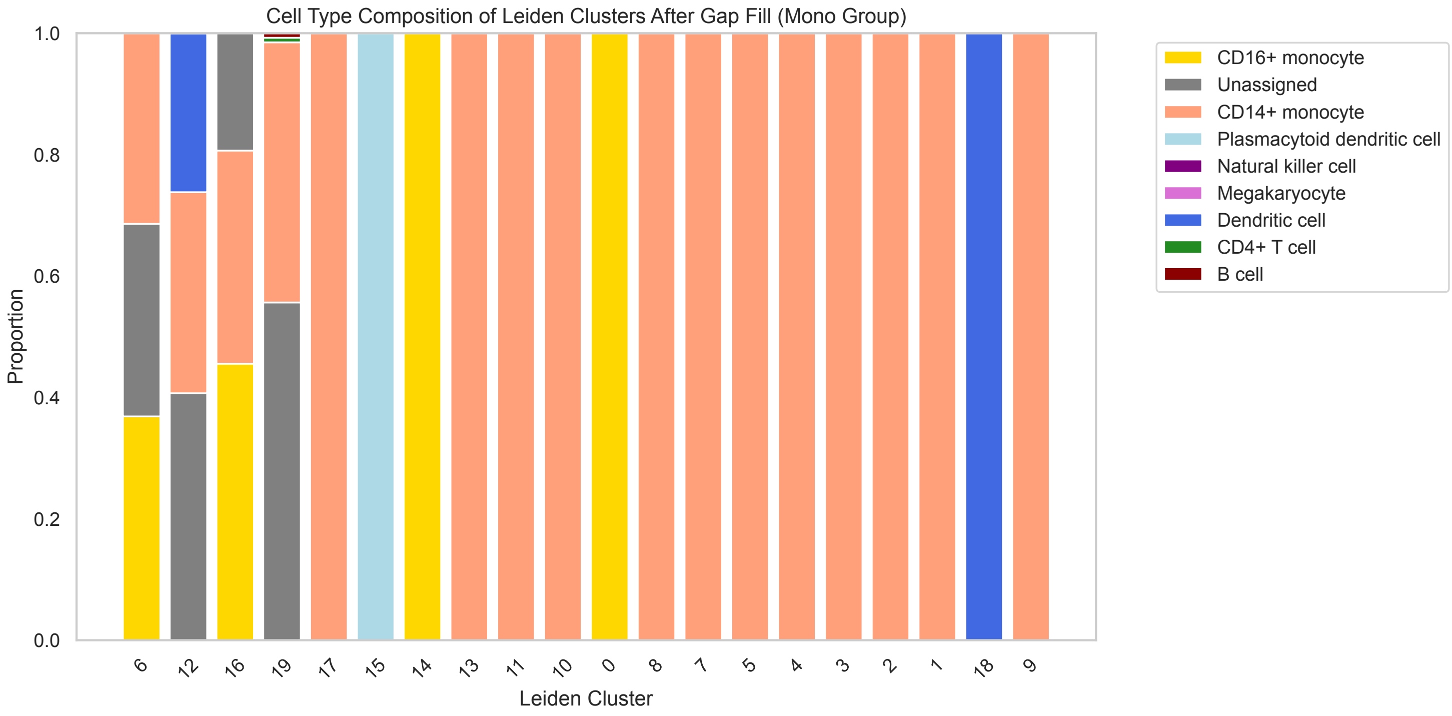


**k**

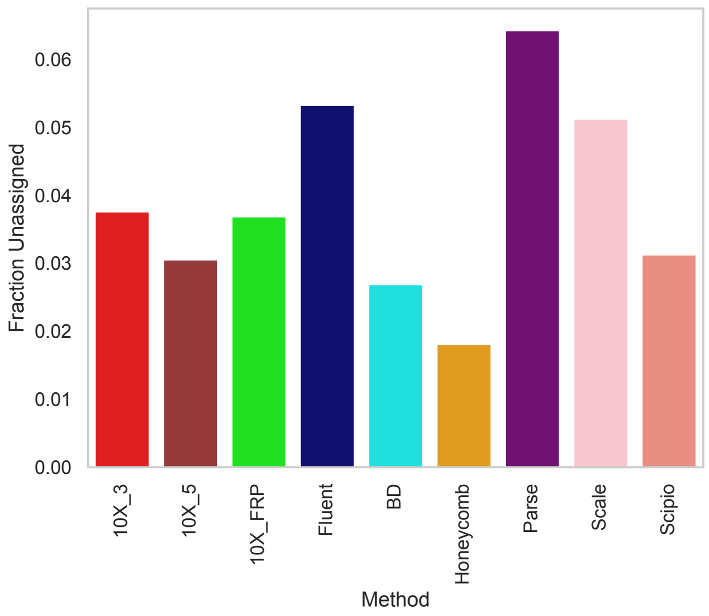


**Extended Data Figure 5**

I

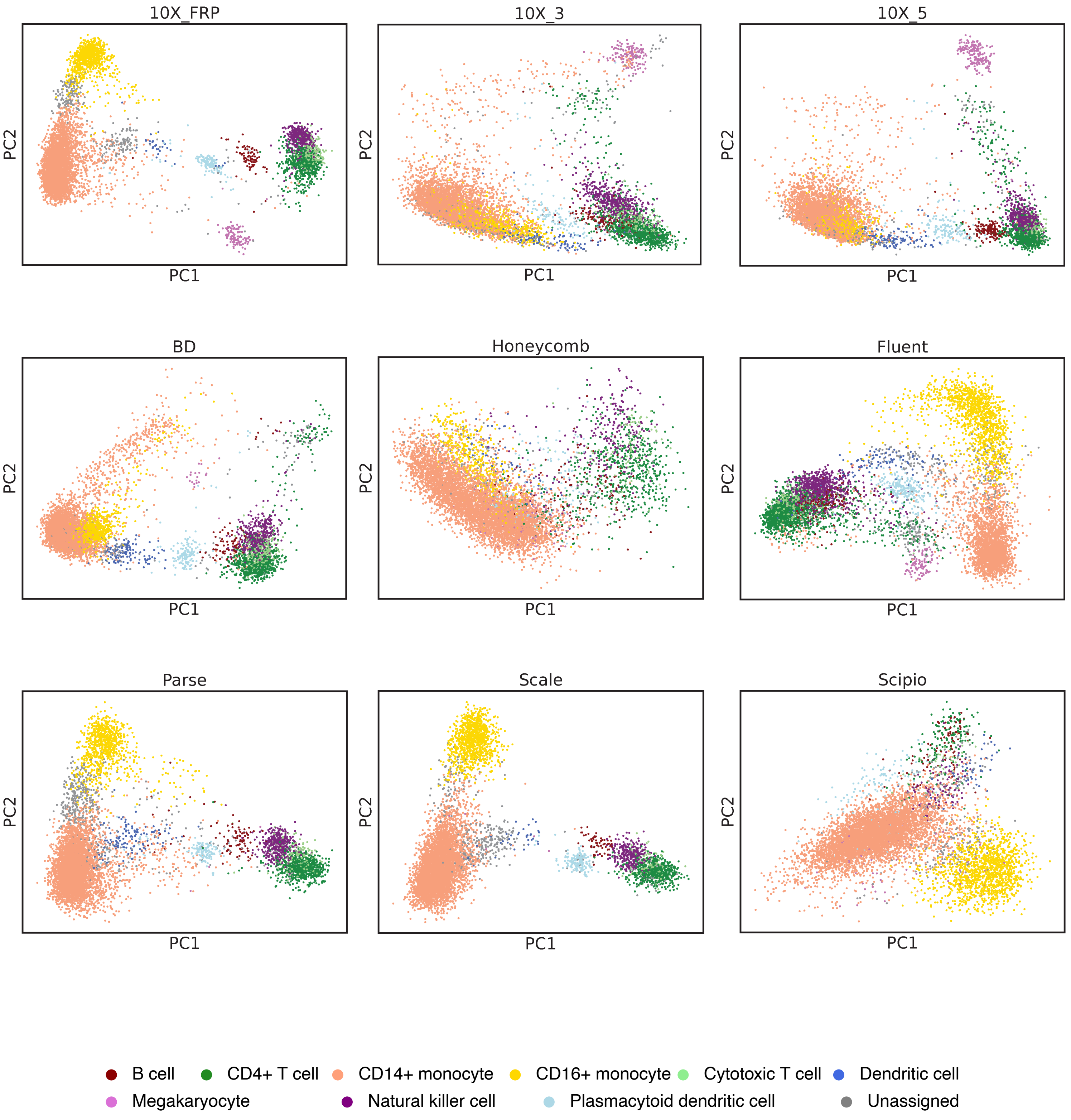


**Extended Data Figure 5**

m

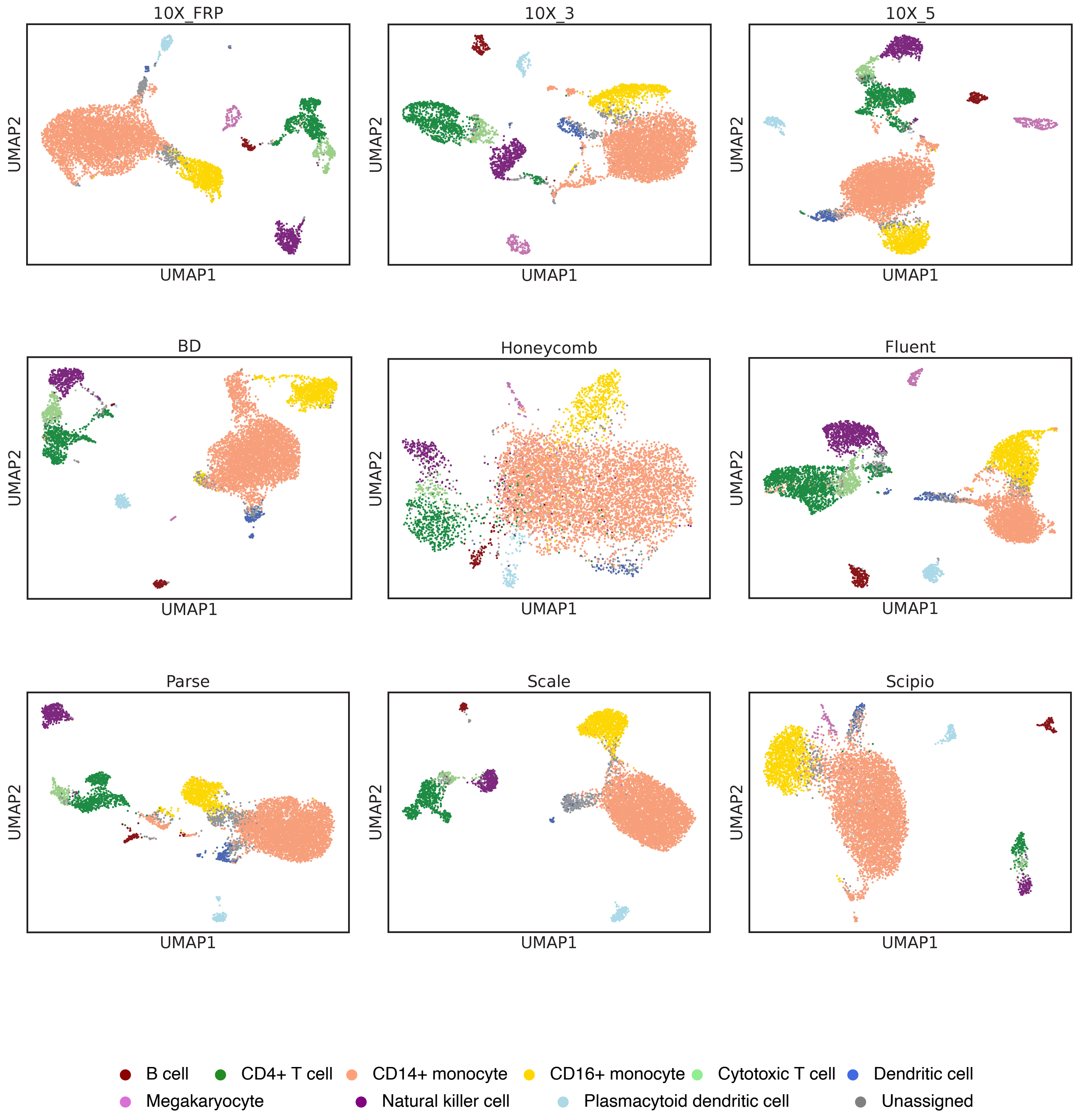


**Extended Data Figure 5**

**a**

### Emulsion-based protocols

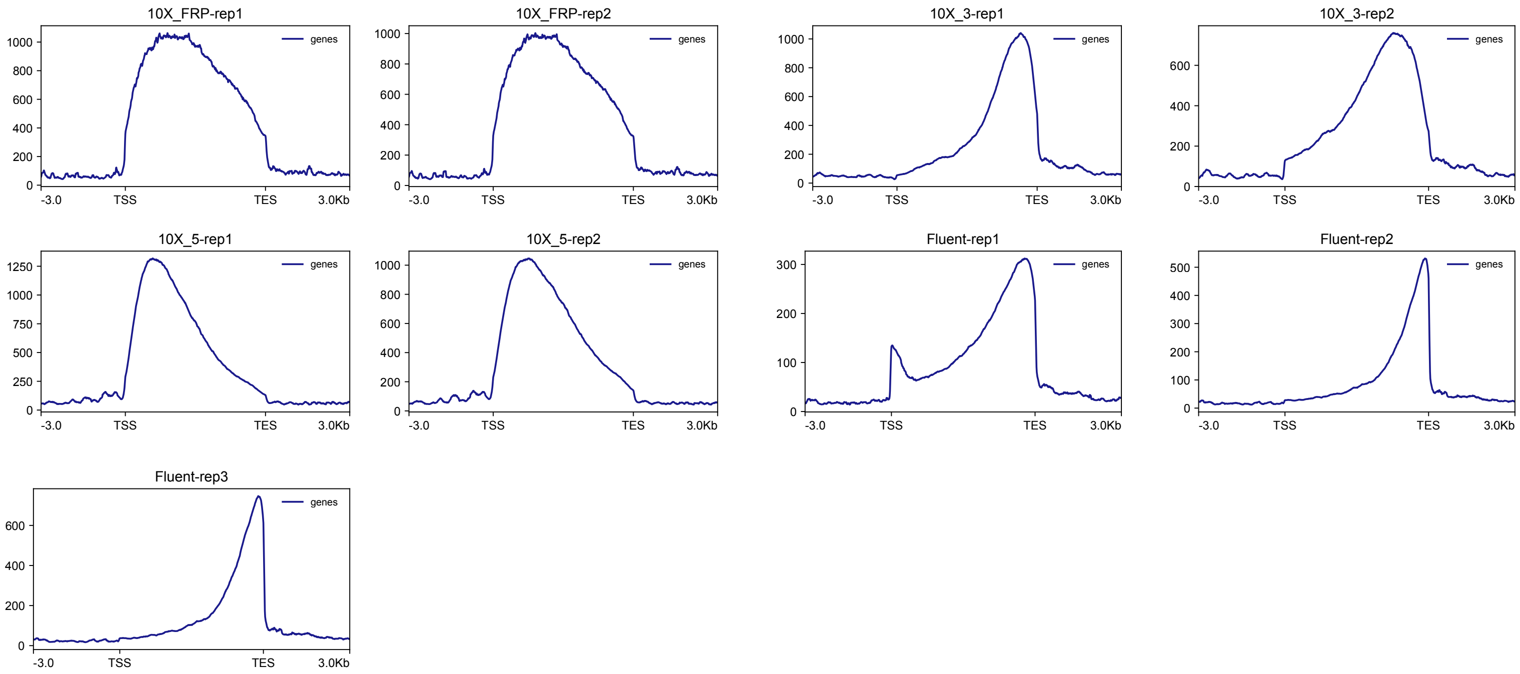


**b**

### Microwell-based protocols

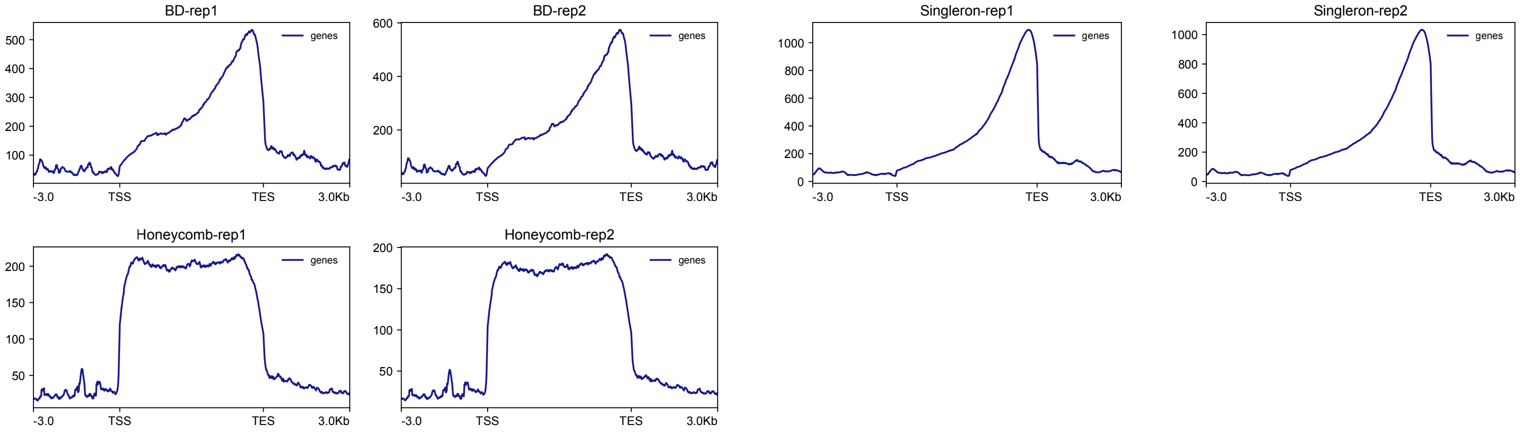


**c**

### Combinatorial indexing protocols

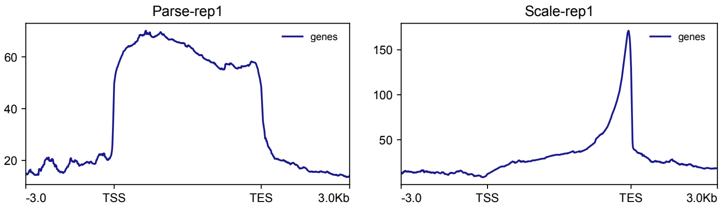


**d**

### Matrigel-based protocols

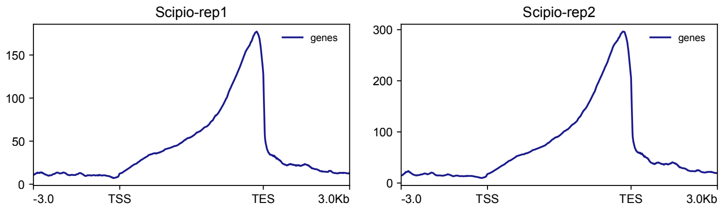


**Extended Data Figure 6**
