## Supplemental Tables for "Comparative Analysis of Commercial Single-Cell RNA Sequencing Technologies"

| kit | cell_recovery_pct |
| --- | --- |
| BD-rep1 | 56.665 |
| Scipio-rep1 | 70.62 |
| 10X_FRP-rep2 | 80.48 |
| 10X_5-rep2 | 81.35 |
| Scale-rep1 | 83.66666667 |
| BD-rep2 | 84.205 |
| 10X_3-rep1 | 84.53 |
| 10X_5-rep1 | 84.55 |
| Fluent-rep3 | 90.52 |
| Scipio-rep2 | 91.16 |
| Honeycomb-rep | 100 |
| Honeycomb-rep | 100 |
| Fluent-rep2 | 103.06 |
| 10X_FRP-rep1 | 103.16 |
| 10X_3-rep2 | 112.68 |
| Fluent-rep1 | 113.33 |
| Parse-rep1 | 122.4561955 |
| Singleron-rep1 | 230.66 |
| Singleron-rep2 | 258.26 |

**Supplementary Table 1a**

| kit | cell_recovery_pct |
| --- | --- |
| 10X_3 | 98.605 |
| 10X_5 | 82.95 |
| 10X_FRP | 91.82 |
| BD | 70.435 |
| Fluent | 102.3033333 |
| Honeycomb | 100 |
| Parse | 122.4561955 |
| Scale | 83.66666667 |
| Scipio | 80.89 |
| Singleron | 244.46 |

**Supplementary Table 1b**

| kit | max_val | RD50 | max_val_sd | RD50_sd | RMSE | R-squared |
|---|---|---|---|---|---|---|
| 10X_FRP | 5335.14395 | 8.206504295 | 12.9368103 | 0.059014518 | 14.01010578 | 0.999754476 |
| 10X_3 | 4184.432458 | 9.800238976 | 19.56554046 | 0.124784129 | 18.23232523 | 0.999309094 |
| BD | 3753.77472 | 9.989917456 | 27.34935358 | 0.195156045 | 24.9098625 | 0.998441575 |
| Parse | 3230.778785 | 11.36430139 | 12.45858744 | 0.110322576 | 6.855523994 | 0.999840754 |
| 10X_5 | 2645.528562 | 8.300568362 | 33.62036054 | 0.311918408 | 36.01611434 | 0.993257856 |
| Scale | 2083.462058 | 5.424652098 | 4.198793748 | 0.040325609 | 4.208099928 | 0.999839228 |
| Fluent | 1890.570885 | 12.75227459 | 184.7225489 | 3.013609914 | 173.8710355 | 0.770302849 |
| Singleron | 1625.36044 | 5.51733888 | 17.08142136 | 0.227748784 | 24.49498084 | 0.988066848 |
| Honeycomb | 1209.738957 | 7.354778182 | 34.76236771 | 0.66091513 | 40.98605453 | 0.96073551 |
| Scipio | 828.6396071 | 4.754690171 | 42.2322821 | 0.941641071 | 67.04596901 | 0.786552601 |

**Supplementary Table 2**

| kit | max_val | RD50 | max_val_sd | RD50_sd | RMSE | R-squared |
|---|---|---|---|---|---|---|
| 10X_FRP | 26410.24362 | 28.18742384 | 205.7764579 | 0.375893607 | 59.80548803 | 0.999724235 |
| BD | 17384.19984 | 27.47179754 | 309.6588876 | 0.844251271 | 92.59357228 | 0.998510108 |
| 10X_3 | 17237.26326 | 26.79029351 | 157.5565921 | 0.427900998 | 49.13445909 | 0.999574509 |
| Parse | 9577.097879 | 28.86537268 | 41.42924819 | 0.211518542 | 7.903095857 | 0.999962805 |
| 10X_5 | 7534.629559 | 14.59392042 | 69.50755963 | 0.302715128 | 43.92431546 | 0.998715873 |
| Singleron | 4642.322905 | 10.80493103 | 96.86419646 | 0.610250298 | 80.9800125 | 0.986713819 |
| Fluent | 4106.38788 | 20.45179892 | 834.3721328 | 8.16057139 | 472.1157336 | 0.655331178 |
| Scale | 3841.408501 | 8.667761701 | 24.57790227 | 0.160120237 | 17.2307482 | 0.999290489 |
| Honeycomb | 2840.24223 | 15.43181921 | 114.7194238 | 1.361407468 | 67.97110008 | 0.978465711 |
| Scipio | 1631.0859 | 7.836993124 | 145.1387189 | 2.086396811 | 160.1235916 | 0.742401826 |

**Supplementary Table 3**

| kit | on_target_mapped | off_target_mapped | unmapped/lost |
|---|---|---|---|
| 10X_FRP-rep1 | 0.947023482 | 0.037044957 | 0.015931561 |
| 10X_FRP-rep2 | 0.900279363 | 0.089236192 | 0.010484445 |
| 10X_3-rep1 | 0.883098324 | 0.084899295 | 0.032002381 |
| 10X_3-rep2 | 0.88107438 | 0.08366062 | 0.035264999 |
| 10X_5-rep1 | 0.791336391 | 0.064008249 | 0.14465536 |
| 10X_5-rep2 | 0.778026635 | 0.059008803 | 0.162964562 |
| BD-rep2 | 0.748106592 | 0.18198107 | 0.069912338 |
| BD-rep1 | 0.713809807 | 0.205526489 | 0.080663704 |
| Scale-rep1 | 0.644009132 | 0.115875644 | 0.240115224 |
| Scipio-rep2 | 0.582008245 | 0.092487709 | 0.325504046 |
| Scipio-rep1 | 0.54916287 | 0.110046106 | 0.340791024 |
| Honeycomb-rep1 | 0.546324287 | 0.326674093 | 0.12700162 |
| Honeycomb-rep2 | 0.496964457 | 0.371761037 | 0.131274507 |
| Singleron-rep1 | 0.486726156 | 0.195566415 | 0.317707429 |
| Singleron-rep2 | 0.474927997 | 0.214472658 | 0.310599344 |
| Parse-rep1 | 0.45982162 | 0.093941659 | 0.446236721 |
| Fluent-rep1 | 0.384645619 | 0.27926472 | 0.336089661 |
| Fluent-rep3 | 0.332533841 | 0.321364127 | 0.346102032 |
| Fluent-rep2 | 0.325877269 | 0.28582728 | 0.388295451 |

**Supplementary Table 4A**

| kit | on_target_mapped | off_target_mapped | unmapped/lost |
|---|---|---|---|
| 10X_3 | 0.882086352 | 0.084279957 | 0.03363369 |
| 10X_5 | 0.784681513 | 0.061508526 | 0.153809961 |
| 10X_FRP | 0.923651422 | 0.063140574 | 0.013208003 |
| BD | 0.7309582 | 0.193753779 | 0.075288021 |
| Fluent | 0.347685576 | 0.295485375 | 0.356829048 |
| Honeycomb | 0.521644372 | 0.349217565 | 0.129138063 |
| Parse | 0.45982162 | 0.093941659 | 0.446236721 |
| Scale | 0.644009132 | 0.115875644 | 0.240115224 |
| Scipio | 0.565585558 | 0.101266907 | 0.333147535 |
| Singleron | 0.480827077 | 0.205019536 | 0.314153387 |

**Supplementary Table 4B**

| kit | total_umi_recovery |
| --- | --- |
| Scipio | 0.054319573 |
| Honeycomb | 0.088145763 |
| Fluent | 0.104514017 |
| Scale | 0.110104916 |
| Singleron | 0.125011209 |
| 10X_5 | 0.18570749 |
| Parse | 0.188088972 |
| BD | 0.304093198 |
| 10X_3 | 0.320607052 |
| 10X_FRP | 0.442232408 |

**Supplementary Table 5**

| kit | n_genes_by_counts | total_umi_counts |
|---|---|---|
| 10X_3-rep1 | 3144 | 8899.5 |
| 10X_3-rep2 | 3136 | 9015.5 |
| 10X_5-rep1 | 2001 | 4950.5 |
| 10X_5-rep2 | 2109 | 4994 |
| 10X_FRP-rep1 | 4282 | 13632 |
| 10X_FRP-rep2 | 4312 | 13719 |
| BD-rep1 | 2850.5 | 9152.5 |
| BD-rep2 | 2793 | 8779 |
| Fluent-rep1 | 1553 | 3142.5 |
| Fluent-rep2 | 1252 | 2065 |
| Fluent-rep3 | 1138 | 1982 |
| Honeycomb-rep1 | 1025 | 1972 |
| Honeycomb-rep2 | 920 | 1778 |
| Parse-rep1 | 2324 | 4791 |
| Scale-rep1 | 1748 | 2937 |
| Scipio-rep1 | 811 | 1522 |
| Scipio-rep2 | 627 | 1063 |

**Supplementary Table 6A**

| kit | n_genes_by_counts | total_umi_counts |
| --- | --- | --- |
| 10X_3 | 3052.03071 | 9460.196065 |
| 10X_5 | 2115.170774 | 5463.866 |
| 10X_FRP | 4071.614516 | 12939.17219 |
| BD | 2754.204581 | 8991.70871 |
| Fluent | 1484.03514 | 3077.075613 |
| Honeycomb | 1188.543548 | 2626.555097 |
| Parse | 2412.950065 | 5465.031097 |
| Scale | 1783.925806 | 3174.515355 |
| Scipio | 786.3401421 | 1534.064578 |

**Supplementary Table 6B**

| kit | 10X_3 | 10X_5 | 10X_FRP | BD | Fluent | Honeycomb | Parse | Scale | Scipio |
|---|---|---|---|---|---|---|---|---|---|
| 10X_3 | 1 | 0 | 0 | 2.53E-26 | 0 | 0 | 1.64E-203 | 0 | 0 |
| 10X_5 | 0 | 1 | 0 | 0 | 0 | 0 | 3.94E-72 | 4.59E-94 | 0 |
| 10X_FRP | 0 | 0 | 1 | 0 | 0 | 0 | 0 | 0 | 0 |
| BD | 2.53E-26 | 0 | 0 | 1 | 0 | 0 | 4.15E-102 | 0 | 0 |
| Fluent | 0 | 0 | 0 | 0 | 1 | 1.43E-179 | 0 | 6.70E-136 | 0 |
| Honeycomb | 0 | 0 | 0 | 0 | 1.43E-179 | 1 | 0 | 0 | 2.86E-192 |
| Parse | 1.64E-203 | 3.94E-72 | 0 | 4.15E-102 | 0 | 0 | 1 | 2.52E-247 | 0 |
| Scale | 0 | 4.59E-94 | 0 | 0 | 6.70E-136 | 0 | 2.52E-247 | 1 | 0 |
| Scipio | 0 | 0 | 0 | 0 | 0 | 2.86E-192 | 0 | 0 | 1 |

**Supplementary Table 7**

| kit | 10X_3 | 10X_5 | 10X_FRP | BD | Fluent | Honeycomb | Parse | Scale | Scipio |
|---|---|---|---|---|---|---|---|---|---|
| 10X_3 | 1 | 0 | 8.24E-254 | 0.03258996 | 0 | 0 | 0 | 0 | 0 |
| 10X_5 | 0 | 1 | 0 | 0 | 0 | 0 | 1 | 0 | 0 |
| 10X_FRP | 8.24E-254 | 0 | 1 | 5.79E-207 | 0 | 0 | 0 | 0 | 0 |
| BD | 0.03258996 | 0 | 5.79E-207 | 1 | 0 | 0 | 0 | 0 | 0 |
| Fluent | 0 | 0 | 0 | 0 | 1 | 1.92E-44 | 0 | 1.27E-21 | 0 |
| Honeycomb | 0 | 0 | 0 | 0 | 1.92E-44 | 1 | 0 | 4.27E-87 | 8.87E-208 |
| Parse | 0 | 1 | 0 | 0 | 0 | 0 | 1 | 3.47E-307 | 0 |
| Scale | 0 | 0 | 0 | 0 | 1.27E-21 | 4.27E-87 | 3.47E-307 | 1 | 0 |
| Scipio | 0 | 0 | 0 | 0 | 0 | 8.87E-208 | 0 | 0 | 1 |

**Supplementary Table 8**

| kit | pct_counts_mt | pct_counts_ribo |
|---|---|---|
| 10X_3-rep1 | 0.033216545 | 0.13586361 |
| 10X_3-rep2 | 0.047405494 | 0.138451807 |
| 10X_5-rep1 | 0.022848325 | 0.190157279 |
| 10X_5-rep2 | 0.019134874 | 0.16594471 |
| 10X_FRP-rep1 | 0.001755359 | 0 |
| 10X_FRP-rep2 | 0.000417032 | 0 |
| BD-rep1 | 0.141056351 | 0.08937213 |
| BD-rep2 | 0.138810351 | 0.085748334 |
| Fluent-rep1 | 0.032215349 | 0.096363585 |
| Fluent-rep2 | 0.035028996 | 0.059934588 |
| Fluent-rep3 | 0.037401846 | 0.067283172 |
| Honeycomb-rep1 | 0.022855431 | 0.097355865 |
| Honeycomb-rep2 | 0.02612027 | 0.102885798 |
| Parse-rep1 | 0.004659394 | 0.0050837 |
| Scale-rep1 | 0.002273696 | 0.005679585 |
| Scipio-rep1 | 0.005892752 | 0.121951215 |
| Scipio-rep2 | 0.007699711 | 0.137032837 |

**Supplementary Table 9A**

| kit | pct_counts_mt | pct_counts_ribo |
|---|---|---|
| 10X_3 | 0.050316729 | 0.160995977 |
| 10X_5 | 0.025896116 | 0.190271163 |
| 10X_FRP | 0.00268012 | 0 |
| BD | 0.145125329 | 0.094396154 |
| Fluent | 0.038441725 | 0.135970101 |
| Honeycomb | 0.025425706 | 0.105017128 |
| Parse | 0.004885083 | 0.005307959 |
| Scale | 0.002516255 | 0.006141771 |
| Scipio | 0.010140706 | 0.135714168 |

**Supplementary Table 9B**

| kit | 10X_3 | 10X_5 | 10X_FRP | BD | Fluent | Honeycomb | Parse | Scale | Scipio |
|---|---|---|---|---|---|---|---|---|---|
| 10X_3 | 1 | 0 | 0 | 0 | 1.70E-69 | 0 | 0 | 0 | 0 |
| 10X_5 | 0 | 1 | 0 | 0 | 0 | 1.83E-16 | 0 | 0 | 0 |
| 10X_FRP | 0 | 0 | 1 | 0 | 0 | 0 | 7.31E-159 | 4.12E-13 | 0 |
| BD | 0 | 0 | 0 | 1 | 0 | 0 | 0 | 0 | 0 |
| Fluent | 1.70E-69 | 0 | 0 | 0 | 1 | 0 | 0 | 0 | 0 |
| Honeycomb | 0 | 1.83E-16 | 0 | 0 | 0 | 1 | 0 | 0 | 0 |
| Parse | 0 | 0 | 7.31E-159 | 0 | 0 | 0 | 1 | 6.07E-61 | 5.11E-46 |
| Scale | 0 | 0 | 4.12E-13 | 0 | 0 | 0 | 6.07E-61 | 1 | 9.59E-213 |
| Scipio | 0 | 0 | 0 | 0 | 0 | 0 | 5.11E-46 | 9.59E-213 | 1 |

**Supplementary Table 10**

| kit | gene_max | total_genes_detected | pct_genes_detected | mt_max | total_mt_detected | pct_mt_detected | ribo_max | total_ribo_detected | pct_ribo_detected |
|---|---|---|---|---|---|---|---|---|---|
| 10X_3-rep1 | 36601 | 28647 | 0.782683533 | 13 | 13 | 1 | 103 | 101 | 0.9805825 |
| 10X_3-rep2 | 36601 | 29156 | 0.796590257 | 13 | 13 | 1 | 103 | 102 | 0.9902913 |
| 10X_5-rep1 | 36601 | 25491 | 0.696456381 | 13 | 13 | 1 | 103 | 101 | 0.9805825 |
| 10X_5-rep2 | 36601 | 25871 | 0.706838611 | 13 | 13 | 1 | 103 | 101 | 0.9805825 |
| 10X_FRP-rep1 | 18082 | 15308 | 0.846587767 | 11 | 11 | 1 | 0 | 0 | |
| 10X_FRP-rep2 | 18082 | 15095 | 0.834808096 | 11 | 11 | 1 | 0 | 0 | |
| BD-rep1 | 30648 | 30648 | 1 | 13 | 13 | 1 | 101 | 101 | 1 |
| BD-rep2 | 30772 | 30772 | 1 | 13 | 13 | 1 | 102 | 102 | 1 |
| Fluent-rep1 | 36601 | 34211 | 0.934701238 | 13 | 13 | 1 | 103 | 102 | 0.9902913 |
| Fluent-rep2 | 36601 | 33542 | 0.916423049 | 13 | 13 | 1 | 103 | 102 | 0.9902913 |
| Fluent-rep3 | 36601 | 32842 | 0.897297888 | 13 | 13 | 1 | 103 | 102 | 0.9902913 |
| Honeycomb-rep1 | 24314 | 24314 | 1 | 13 | 13 | 1 | 101 | 101 | 1 |
| Honeycomb-rep2 | 26146 | 26146 | 1 | 13 | 13 | 1 | 102 | 102 | 1 |
| Parse-rep1 | 36601 | 29203 | 0.797874375 | 13 | 13 | 1 | 103 | 100 | 0.9708738 |
| Scale-rep1 | 36601 | 33342 | 0.910958717 | 13 | 13 | 1 | 103 | 101 | 0.9805825 |
| Scipio-rep1 | 36601 | 20377 | 0.556733423 | 13 | 13 | 1 | 103 | 98 | 0.9514563 |
| Scipio-rep2 | 36601 | 20110 | 0.54943854 | 13 | 13 | 1 | 103 | 99 | 0.961165 |

**Supplementary Table 11**

| kit | 10X_3 | 10X_5 | 10X_FRP | BD | Fluent | Honeycomb | Parse | Scale | Scipio |
|---|---|---|---|---|---|---|---|---|---|
| 10X_3 | 1 | 9.01E-160 | 0 | 0 | 0 | 0 | 0 | 0 | 6.39E-09 |
| 10X_5 | 9.01E-160 | 1 | 0 | 0 | 0 | 0 | 0 | 0 | 2.22E-178 |
| 10X_FRP | 0 | 0 | 1 | 0 | 0 | 0 | 1.12E-200 | 3.26E-242 | 0 |
| BD | 0 | 0 | 0 | 1 | 8.01E-51 | 2.27E-50 | 0 | 0 | 0 |
| Fluent | 0 | 0 | 0 | 8.01E-51 | 1 | 1 | 0 | 0 | 1.51E-200 |
| Honeycomb | 0 | 0 | 0 | 2.27E-50 | 1 | 1 | 0 | 0 | 1.85E-165 |
| Parse | 0 | 0 | 1.12E-200 | 0 | 0 | 0 | 1 | 0.336930772 | 0 |
| Scale | 0 | 0 | 3.26E-242 | 0 | 0 | 0 | 0.336930772 | 1 | 0 |
| Scipio | 6.39E-09 | 2.22E-178 | 0 | 0 | 1.51E-200 | 1.85E-165 | 0 | 0 | 1 |

**Supplementary Table 12**

| kit | count |
| --- | --- |
| 10X_3-rep1 | 28 |
| 10X_3-rep2 | 31 |
| 10X_5-rep1 | 18 |
| 10X_5-rep2 | 23 |
| 10X_FRP-rep1 | 11 |
| 10X_FRP-rep2 | 16 |
| BD-rep1 | 42 |
| BD-rep2 | 23 |
| Fluent-rep1 | 358 |
| Fluent-rep2 | 259 |
| Fluent-rep3 | 151 |
| Honeycomb-rep1 | 15 |
| Honeycomb-rep2 | 33 |
| Parse-rep1 | 70 |
| Scale-rep1 | 342 |
| Scipio-rep1 | 2 |
| Scipio-rep2 | 1 |

**Supplementary Table 13**

| feature | pc1 | pc2 |
|---|---|---|
| gene_length | 0.68 | 0.12 |
| gene_GC | -0.41 | -0.02 |
| MT_score | 0.02 | -0.42 |
| total_UMI | -0.13 | 0.6 |

**Supplementary Table 14**

| CellType | precision | recall | f1-score | support |
|---|---|---|---|---|
| B cell | 0.996713229 | 0.973515249 | 0.98497767 | 1246 |
| CD14+ monocyte | 0.935205184 | 0.944727273 | 0.939942113 | 1375 |
| CD16+ monocyte | 0.909090909 | 0.663265306 | 0.766961652 | 196 |
| CD4+ T cell | 0.932087227 | 0.79321315 | 0.857061014 | 1886 |
| Cytotoxic T cell | 0.923119777 | 0.737099644 | 0.81968835 | 2248 |
| Natural killer cell | 0.928205128 | 0.480106101 | 0.632867133 | 377 |
| Plasmacytoid dendritic cell | 0.866666667 | 0.702702703 | 0.776119403 | 37 |
| Dendritic cell | 1 | 0.236842105 | 0.382978723 | 114 |
| Megakaryocyte | 0.995951417 | 0.968503937 | 0.982035928 | 254 |
| Unassigned | 0.001814882 | 0.117647059 | 0.00357462 | 17 |
| accuracy | | | 0.809935484 | 7750 |
| macro avg | 0.848885442 | 0.661762253 | 0.714620661 | 7750 |
| weighted avg | 0.940398182 | 0.809935484 | 0.863170109 | 7750 |

**Supplementary Table 15**

| kit | bARI | bAMI | ARI | AMI | ASW | CVR10 | hmean |
|---|---|---|---|---|---|---|---|
| 10X_3 | 0.744673953 | 0.839356767 | 0.847407942 | 0.810831812 | 0.618140861 | 0.2844578 | 0.595188329 |
| 10X_5 | 0.812517112 | 0.8546014 | 0.886564434 | 0.828389276 | 0.619937122 | 0.20574734 | 0.53526562 |
| 10X_FRP | 0.726481256 | 0.833632749 | 0.526602619 | 0.730477302 | 0.678899124 | 0.28776595 | 0.555966393 |
| BD | 0.701595591 | 0.80124068 | 0.780426462 | 0.761382017 | 0.621081077 | 0.21207237 | 0.517526643 |
| Fluent | 0.718775754 | 0.816148144 | 0.782259127 | 0.78957474 | 0.618686967 | 0.15448049 | 0.452795633 |
| Honeycomb | 0.454597373 | 0.601597394 | 0.259807913 | 0.48621399 | 0.558142744 | 0.062840275 | 0.218398527 |
| Parse | 0.65167007 | 0.773233401 | 0.475991898 | 0.665291929 | 0.58915703 | 0.14072543 | 0.393826336 |
| Scale | 0.606611488 | 0.740170052 | 0.437309698 | 0.694595073 | 0.617039926 | 0.109715626 | 0.343621912 |
| Scipio | 0.530495869 | 0.72691785 | 0.343288667 | 0.572156729 | 0.569045834 | 0.05502165 | 0.215413056 |

**Supplementary Table 16**

| kit | count | mean | std | min | 25% | 50% | 75% | max |
|---|---|---|---|---|---|---|---|---|
| 10x-3' | 9 | 2259.722222 | 1375.657164 | 1000.5 | 1402 | 1660.5 | 2413.5 | 5509.5 |
| 10x-5' | 9 | 1478.611111 | 921.0773847 | 834.5 | 921 | 1084.5 | 1553 | 3733 |
| 10x-FRP | 9 | 2782.777778 | 1493.29983 | 1208.5 | 2093 | 2679.5 | 2868.5 | 6437 |
| Fluent | 9 | 1090.666667 | 594.3790878 | 481 | 722.3333333 | 946.3333333 | 1313.666667 | 2245 |
| BD | 9 | 1305.388889 | 466.9544128 | 764.5 | 952 | 1185.5 | 1821.5 | 2045.5 |
| Honeycomb | 9 | 252.9444444 | 130.8490171 | 90 | 186 | 206 | 291 | 521 |
| Parse | 8 | 917.875 | 431.482308 | 474 | 579.25 | 828.5 | 1170.75 | 1608 |
| Scale | 8 | 670.375 | 393.179689 | 61 | 420 | 631.5 | 952.75 | 1234 |
| Scipio | 9 | 213.0555556 | 108.2821674 | 65 | 142.5 | 204.5 | 284.5 | 365.5 |

**Supplementary Table 17**

| kit | slope | constant | n=10^4 | n=10^5 | n=10^6 | n=10^7 |
|---|---|---|---|---|---|---|
| 10X_3p | 0.16 | 65000 | 66600 | 81000 | 225000 | 1665000 |
| 10X_5p | 0.16 | 65000 | 66600 | 81000 | 225000 | 1665000 |
| 10X_FRP | 0.17 | 65000 | 66700 | 82000 | 235000 | 1765000 |
| Fluent | 0.05 | 3000 | 3500 | 8000 | 53000 | 503000 |
| BD | 0.08 | 11000 | 11800 | 19000 | 91000 | 811000 |
| Singleron | 0.05 | 60000 | 60500 | 65000 | 110000 | 560000 |
| Honeycomb | 0.11 | 2000 | 3100 | 13000 | 112000 | 1102000 |
| Scipio | 0.21 | 0 | 2100 | 21000 | 210000 | 2100000 |
| Parse | 0.11 | 0 | 1100 | 11000 | 110000 | 1100000 |
| Scale | 0.09 | 0 | 900 | 9000 | 90000 | 900000 |

**Supplementary Table 18A**

| kit | time | n_cells | n=10^4 | n=10^5 | n=10^6 | n=10^7 | rate | factor |
|---|---|---|---|---|---|---|---|---|
| 10X_3p | 8.5 | 10000 | 8.5 | 85 | 850 | 8500 | 1176.470588 | 2.882352941 |
| 10X_5p | 8.5 | 10000 | 8.5 | 85 | 850 | 8500 | 1176.470588 | 2.882352941 |
| 10X_FRP | 24.5 | 10000 | 24.5 | 245 | 2450 | 24500 | 408.1632653 | 1 |
| Fluent | 14.5 | 20000 | 7.25 | 72.5 | 725 | 7250 | 1379.310345 | 3.379310345 |
| BD | 7.83 | 20000 | 3.915 | 39.15 | 391.5 | 3915 | 2554.278416 | 6.25798212 |
| Singleron | 7.5 | 10000 | 7.5 | 75 | 750 | 7500 | 1333.333333 | 3.266666667 |
| Honeycomb | 10.5 | 17000 | 6.176470588 | 61.76470588 | 617.6470588 | 6176.470588 | 1619.047619 | 3.966666667 |
| Scipio | 11.33 | 5000 | 22.66 | 226.6 | 2266 | 22660 | 441.3062665 | 1.081200353 |
| Parse | 19 | 140000 | 1.357142857 | 13.57142857 | 135.7142857 | 1357.142857 | 7368.421053 | 18.05263158 |
| Scale | 10.33 | 153600 | 0.672526042 | 6.725260417 | 67.25260417 | 672.5260417 | 14869.31268 | 36.42981607 |

**Supplementary Table 18B**

| kit | doublet_rate |
|---|---|
| 10X_3-rep1 | 0.032887732 |
| 10X_3-rep2 | 0.02715655 |
| 10X_5-rep1 | 0.032879953 |
| 10X_5-rep2 | 0.028764597 |
| 10X_FRP-rep1 | 0.036545173 |
| 10X_FRP-rep2 | 0.037027833 |
| BD-rep1 | 0.024177182 |
| BD-rep2 | 0.033192803 |
| Fluent-rep1 | 0.008294362 |
| Fluent-rep2 | 0.007665438 |
| Fluent-rep3 | 0.005302696 |
| Honeycomb-rep1 | 0.03160316 |
| Honeycomb-rep2 | 0.0247 |
| Singleron-rep1 | 0.016777942 |
| Singleron-rep2 | 0.021761016 |
| Scipio-rep1 | 0.020957236 |
| Scipio-rep2 | 0.029179465 |
| Parse-rep1 | 0.028553096 |
| Scale-rep1 | 0.034611554 |
| Broad-Reference-Val | 0.0112 |
| Broad-Reference | 0.008610437 |

**Supplementary Table 19**

| Isotope | Element | Target | Clone |
|---|---|---|---|
| 89 | Y | CD45 | HI30 |
| 110 | Cd | CD117 | 104D2 |
| 111 | Cd | CD10 | HI10a |
| 112 | Cd | CD11b | ICRF44 |
| 113 | Cd | CD7 | 6B7 |
| 114 | Cd | CD1c | L161 |
| 115 | In | CD57 | HCD57 |
| 116 | Cd | CD33 | WM53 |
| 141 | Pr | 9d (alpha 4 inte | 9F10 |
| 143 | Nd | CD127 | A019D5 |
| 144 | Nd | CD15 | W6D3 |
| 145 | Nd | CD4 | RPA-T4 |
| 146 | Nd | IgD | 1A6-2 |
| 149 | Sm | CD177 | MEM-166 |
| 150 | Nd | amma delta TCI | Sa6-E9 |
| 151 | Eu | CD123 | 6H6 |
| 154 | Sm | CD3 | UCHT1 |
| 155 | Gd | CD27 | L128 |
| 156 | Gd | CD39 | A1 |
| 157 | Gd | CD19 | HIB19 |
| 158 | Gd | Vα7.2 | 3C10 |
| 159 | Tb | CD11c | Bu15 |
| 160 | Gd | CD56 | REA196 |
| 161 | Dy | CD66 | B1.1 |
| 163 | Dy | CD20 | 2H7 |
| 164 | Dy | CD161 | HP-3G10 |
| 165 | Ho | CD45RO | UCHL1 |
| 166 | Er | FcERI | AER-37 |
| 167 | Er | CCR7 | G043H7 |
| 168 | Er | CD8 | SK1 |
| 169 | Tm | CD25 | 2A3 |
| 170 | Er | CD45RA | HI100 |
| 172 | Yb | CD14 | M5E2 |
| 174 | Yb | HLADR | L243 |
| 175 | Lu | PD1 | EH12.2H7 |
| 176 | Yb | CD38 | HIT2 |
| 209 | Bi | CD16 | 3G8 |

**Supplementary Table 20A**

| Population | Gating Strategy | % of Live Cells |
|---|---|---|
| CD3+ T Cells | Live/CD45+/ CD3+ | 8.28 |
| CD4+ T cells | Live/CD45+/ CD3+/CD4+ | 4.8 |
| CD8+ T Cells | Live/CD45+/ CD3+/CD8+ | 3.09 |
| gd T Cells | Live/CD45+/ CD3+/ gamma delta TCR+ | 0.92 |
| B Cells | Live/CD45+/ CD3-/ CD19+ | 1.04 |
| NK Cells | Live/CD45+/ CD3-/ CD19-/CD56+ | 5.63 |
| CD14 Mono | Live/CD45+/ CD3-/ CD19-/CD56-/CD11c+/HLADR+/CD14+/CD16- | 50.7 |
| CD16 Mono | Live/CD45+/ CD3-/ CD19-/CD56-/CD11c+/HLADR+/CD14-/CD16+ | 6.03 |
| Int Mono | Live/CD45+/ CD3-/ CD19-/CD56-/CD11c+/HLADR+/CD14+/CD16+ | 2.44 |
| mDC | Live/CD45+/ CD3-/ CD19-/CD56-/CD11c+/HLADR+/CD16-/CD14-/CD1c | 2.25 |
| pDC | Live/CD45+/ CD3-/ CD19-/CD56-/CD11c low/HLADR+/CD123+ | 0.85 |
| Basophils | Live/CD45+/ CD3-/ CD19-/CD56-/CD11c+/HLADR low/CD123+ | 0.59 |
| Eosinophils | Live/CD45-/ CD66+/CD49d | 0.12 |
| Neutrophils | Live/CD45-/ CD66+/CD16+ | 0.1 |

**Supplementary Table 20B**
